# Evolutionary rise of a synaptic mechanism for creating and diversifying key reinforcement signals

**DOI:** 10.64898/2026.07.31.742134

**Authors:** Natalia Rodriguez-Sosa, Lupita Rios, You-Hsin Lin, Volodymyr Rybalchenko, Yash Sharma, Alaa Hajeissa, Ian Chambers, Nien-Shao Wang, Raghav Rajesh, Vishal Narla, Steven J. Shabel

**Author notes:** Corresponding author: Steven Shabel. equal contribution (Natalia Rodriguez-Sosa, Lupita Rios).

## Abstract

Most neurons release either excitatory or inhibitory neurotransmitters. However, multiple inputs to the lateral habenula (LHb) co-transmit glutamate and GABA, transmitters with opposing effects on LHb output. Although the LHb has an established role in reinforcement learning, the adaptive significance of glutamate/GABA co-release remains unclear. Using experimentally informed simulations, we show that GABA co-release is sufficient to produce temporal difference (TD)-like transformations of input activity, computations commonly used for reinforcement learning and behavioral optimization. Heterogeneous GABA-to-glutamate ratios, like those found among LHb neurons ex vivo, produce diverse TD-like computations linked to higher-order decision-making. Single-cell RNA-sequencing analysis and machine-learning image analysis further indicate that glutamate/GABA co-release expanded across vertebrate evolution, from fish to mice, rats, and monkeys. Evolutionary expansion of glutamate/GABA co-release may have supported increasingly sophisticated learning and decision-making that contribute to intelligent behavior.

## Introduction

Neural communication is commonly segregated into opposing modes - excitation and inhibition – and most neurons release neurotransmitters that either promote or suppress the activity of their targets^1^. However, this rule is broken in the rodent lateral habenula (LHb), where multiple inputs co-transmit GABA with glutamate^1–8^ – transmitters with antagonistic actions on membrane potential and firing probability (including in LHb neurons^2,9^). This unusual form of synaptic signaling suggests that LHb neurons integrate excitatory and inhibitory drive in a fundamentally distinct manner from other brain regions.

The LHb occupies a pivotal position between forebrain structures and midbrain monoaminergic centers^10–12^. LHb neurons encode TD-like (i.e., temporal-difference- or derivative-like) value reinforcement signals, which are routed topographically to midbrain dopamine neurons through an inhibitory intermediary^13–21^. Disruption of habenula function increases the activity of midbrain dopamine neurons^22^, diminishes their reinforcement-related error signals^22^, and eliminates flexible decision-making during probability and delay discounting tasks in rats^20^, indicating a critical role for the LHb in reinforcement learning and higher-order decision-making. Habenular TD-like reinforcement signals are driven by the major input from the basal ganglia (entopeduncular nucleus, EPN; homologous to the globus pallidus internus in primates) which sends reward- and aversion-related information to the LHb^23–27^. This input is composed of somatostatin-expressing neurons (SST+; > 90% of all LHb-projecting EPN neurons^23,28^), which co-release glutamate and GABA from individual synaptic vesicles^2,29^. Thus, glutamate/GABA co-releasing inputs to the LHb are poised to impact habenular and dopaminergic signals that contribute to flexible, higher-order decision-making^30–40^.

Despite these advances, the adaptive significance of glutamate/GABA co-release in the LHb remains unclear. How might glutamate/GABA co-release shape computations underlying reinforcement learning? Is glutamate/GABA co-release an ancient feature of habenular circuits, or did it expand during evolution in parallel with more sophisticated learning capacities? Here, we use experimentally informed neural simulations to show that glutamate/GABA co-release is sufficient to produce and diversify TD-like computations. We also use cross-species, machine learning classification of synaptic terminals and analysis of scRNA-seq datasets to identify a pattern consistent with evolutionary expansion of glutamate/GABA co-transmitting terminals in mammalian LHb circuits. These findings indicate that glutamate/GABA co-transmission may have contributed to the evolution of higher-order learning and decision-making.

## Results

### Co-release of GABA with glutamate is sufficient for TD-like computations

To investigate the function of GABA co-release with glutamate in the LHb, we simulated activity in a simple, biophysically realistic output neuron while varying the ratio of GABA (GABA-A)-to-glutamate (AMPA) peak conductance from its inputs (Fig. 1c; see Methods; glutamate/GABA co-releasing inputs have little NMDA transmission^41^). Unlike artificial highly synchronous input stimulation in brain slices^2^, each input fired randomly and independently around a mean firing rate (same for all inputs) that varied between 5 and 160 Hz in different simulations (“tonic” input activity simulations; Fig. 1e,f). The kinetics of glutamatergic and GABAergic transmission were matched to experimental data from stimulation of glutamate/GABA co-releasing basal ganglia inputs to the LHb (Fig. 1b, d). To determine the range of GABA-glutamate transmission ratios for the simulations, we examined the range of GABA-A-R to AMPA-R peak conductance ratios from EPN inputs in mice and rats (Fig. 1a). Both species had, on average, near equal GABA-A-R and AMPA-R peak conductances, although the range was larger and significantly more variable in rats (Fig. 1a; Mann-Whitney, P = 0.39, N = 35 cells from 7 mice and 36 cells from 8 rats; F test to compare variances, F = 3.5, P = .0004; stimulation of SST+ EPN inputs^28^ resulted in a similar average ratio to non-specific EPN input stimulation in mice, N = 12 cells from 5 SST-cre mice, Mann-Whitney, P = .28, Fig. S1a). Importantly, variation in GABA-glutamate ratios among LHb neurons is not random, because different co-releasing inputs onto the same LHb neuron have more similar GABA-glutamate transmission ratios than inputs onto different LHb neurons^29^. Increasing levels of GABA co-release in the simulations decreased spike output, as expected, and also flattened the input-output curve at moderate GABA-glutamate ratios (Fig. 1e). The flattening, and eventual reversal, of the input-output curve was caused by GABA having an asymmetrically larger impact when inputs were more active (Fig. 1f).

**Figure 1.**
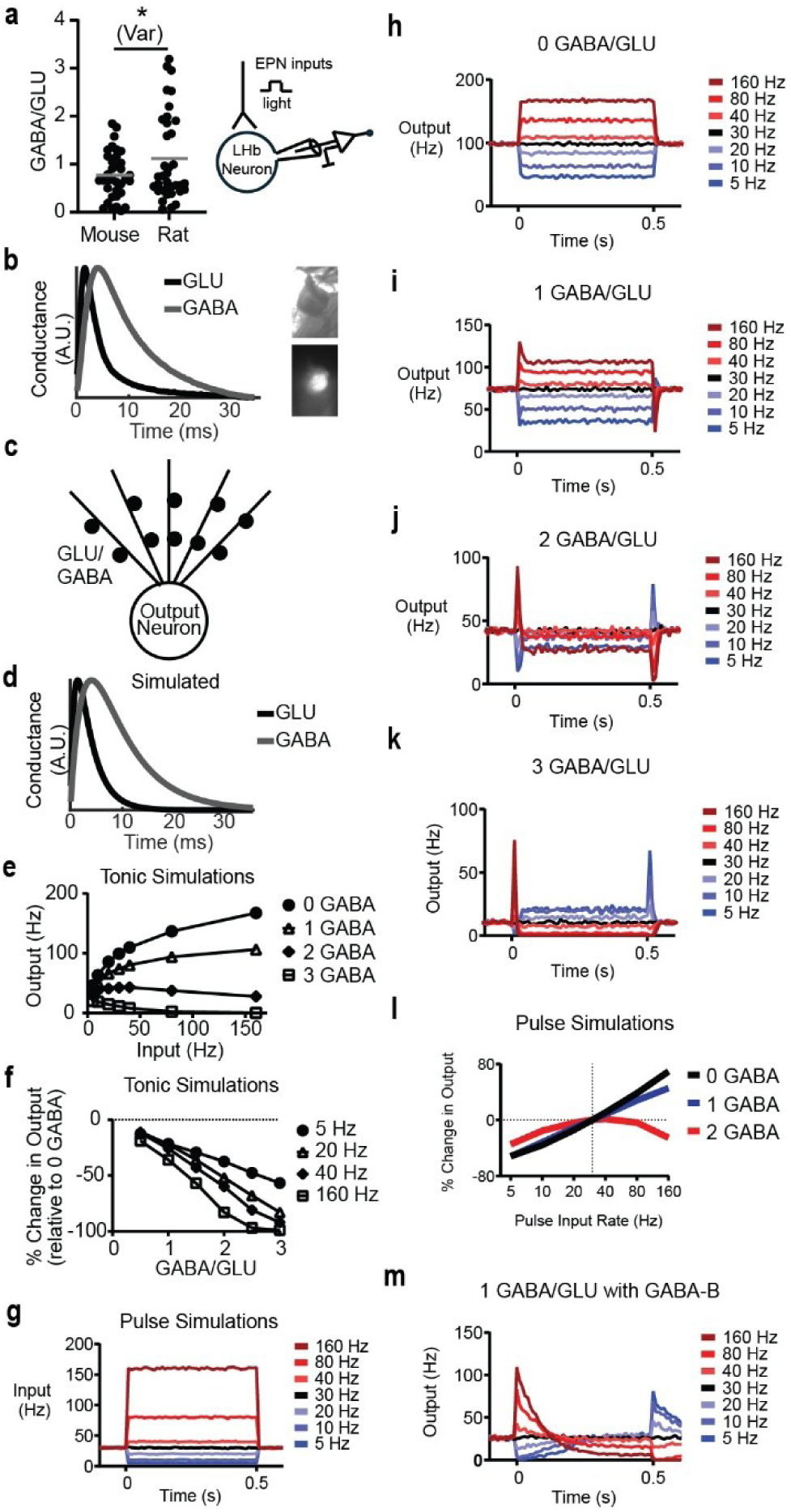
GABA co-release with glutamate as a mechanism for diverse, TD-like computations. **a**, Ratio of peak amplitude of GABA-A-R and AMPA-R-mediated (GLU) currents from whole-cell, ex vivo recordings of mouse and rat LHb neurons (∼ 23 °C) during optogenetic stimulation of EPN inputs. *, P < .005 for comparison of variability. **b**, Normalized AMPA-R and GABA-A-R-mediated currents from whole-cell, ex vivo recordings of mouse LHb neurons (N = 12 neurons from 5 mice; 37 °C) during optogenetic SST+ (glutamate/GABA co-releasing) EPN input stimulation, showing mean kinetics. **c**, Schematic figure of a neuron used for simulations. Dark circles represent synapses from different input neurons. **d**, Kinetics of simulated glutamatergic and GABAergic currents, intended to match those in **b**. **e**, Input-output curves for simulations of tonic input activity with different ratios of GABAergic-to-glutamatergic peak conductances (average of 10, 3-second simulations). GABAergic peak conductance was varied relative to constant glutamatergic peak conductance in all figures. Note that “1 GABA” = 1 GABA/GLU, etc. **f**, Data from same simulations as **e**, but re-calculated and plotted to show the change in output activity for different amounts of GABA co-transmission (relative to no GABA) and different input activity. **g**, Input activity during pulse input simulations (average of 1000 simulations). Colors indicate mean firing rate of inputs during the 0.5 s pulse period for **g**-**k, m**. **h**-**k**, Output activity for different amounts of GABA co-transmission, as indicated. **l**, Change in output activity during first 200 ms of the pulse relative to the 30 Hz input baseline period for different pulse input rates. There were substantial floor effects in the “3 GABA” condition. **m**, same as **i** but including GABA-B conductance.

The flattening of the input-output curve in the tonic input activity simulations suggested that there are conditions in which LHb neurons are insensitive to changes in the activity of glutamate/GABA co-releasing inputs. However, basal ganglia inputs to the LHb, as well as LHb neurons, are known to convey valence via sudden increases and decreases in activity during presentation of aversive and rewarding stimuli, respectively^13,21,23–27,42–44^. Therefore, we next investigated how GABA co-release affects output during changes in input activity. We simulated changes in activity from inputs with high basal firing rates (∼ 30 Hz), like those found in LHb-projecting, basal ganglia neurons^24^ (Fig. 1g). We used 500 ms duration changes in activity (“pulse simulations”) to allow visualization of how GABA co-release changes output activity in both directions from different starting values. These simulations found that GABA co-release caused output to change briefly and in opposite directions during increases and decreases in activity (Fig. 1g-k). Increases in input activity at pulse onset caused a short-lasting increase in output that rapidly returned toward baseline (< 50 ms), even though input activity was elevated for 500 ms (e.g., Fig. 1j). Decreases in input activity caused brief, phasic decreases in output when inputs co-released GABA (e.g., Fig. 1j-k). Short-lasting changes in output also occurred during the offset of the change in input activity – return from low input activity back to basal levels caused phasic increases in output; return from high input activity back to basal levels caused phasic decreases in output (Fig. 1i- k). Thus, GABA co-release causes the response of the output neuron to resemble the derivative of the input activity (i.e., be “TD-like”). Importantly, differentiation (computing the rate of change, derivative, or TD) is a common component of behavioral optimization algorithms^45^, including TD reinforcement learning^46–48^, a type of learning that the LHb and downstream dopamine circuitry are critically involved in^48–54^.

Notably, the amount of differentiation and temporal shape of the output response varied for different GABA-glutamate ratios (Fig. 1h-k). Low levels of GABA co-release caused only fractional differentiation, which can also be achieved with spike-frequency adaptation mechanisms at slower timescales^55^. However, higher levels of GABA co-release caused full differentiation, and even a delayed reversal of activity. Thus, output neurons with “intentionally” different GABA-glutamate ratios, like those found in the LHb^29^, have diverse levels of differentiation and temporal transformations of input activity. Furthermore, the changes in output for increases versus decreases in input activity (as levels of GABA co-release increased) were asymmetric (Fig. 1l), suggesting that GABA co-release could contribute to the asymmetric, biased reinforcement signals in downstream dopamine neurons that allow learning distributions of reward probability^31,34,35,56^. We note that although these simulations show that GABA co-release with glutamate is sufficient for derivative-like computations, there are likely many factors that affect the GABA-glutamate threshold at which full differentiation occurs, such as GABA-B-related conductances^57–59^ (Fig. 1m), the degree of input synchronization, and the activity of other types of inputs. We also found a small change in results when using simulations with twice as many inputs (Fig. S1b-j).

### Slower GABA kinetics drive TD-like computations

To determine the mechanism by which GABA co-release causes differentiation, we computed the conductance of synaptic GABA (GABA-A-R) and glutamate (AMPA-R) channels in the pulse input simulations. Although changes in the activity of co-releasing inputs caused changes in the conductance of both GABAergic and glutamatergic channels (Fig. S2a-b), GABAergic channels were slower to reach steady-state levels due to their slower kinetics (Fig. S2c-f). This caused the glutamate/GABA conductance ratio to change phasically during the onset and offset of the change in input activity (Fig. 2a-c), similar to changes in spiking of the output neuron in our simulations (Fig. 1i-k; Fig. S1f-h), suggesting that differences in transmission kinetics are responsible for differentiation. Consistent with this prediction, when glutamatergic kinetics were artificially slowed so that they matched GABAergic kinetics (Fig. 2d-i; and vice versa, Fig. S3), differentiation during pulse onset and offset was abolished (Fig. 2g-h; Fig. S3d-e). Thus, differences in glutamatergic and GABAergic transmission kinetics are critical for the ability of GABA co-release to cause differentiation. Equalizing glutamatergic and GABAergic kinetics also caused sudden, catastrophic decreases in output activity during tonic input activity simulations with high levels of GABA (Fig. 2d-e; Fig. S3a-b). However, the asymmetric effects of GABA co-release on output during increases versus decreases in pulse input activity were preserved (Fig. 2i; Fig. S3f), indicating that this effect is due to differences in the reversal potentials of the current flowing through the channels. We note that although we modeled glutamate/GABA co-transmission as occurring from individual release sites due to previous electrophysiological and electron microscopy data^2,29^, this is not required for TD-like computations in our simulations. This is because synchronized glutamate and GABA release from adjacent release sites would also produce phasic changes in the glutamate/GABA conductance ratio.

**Figure 2.**
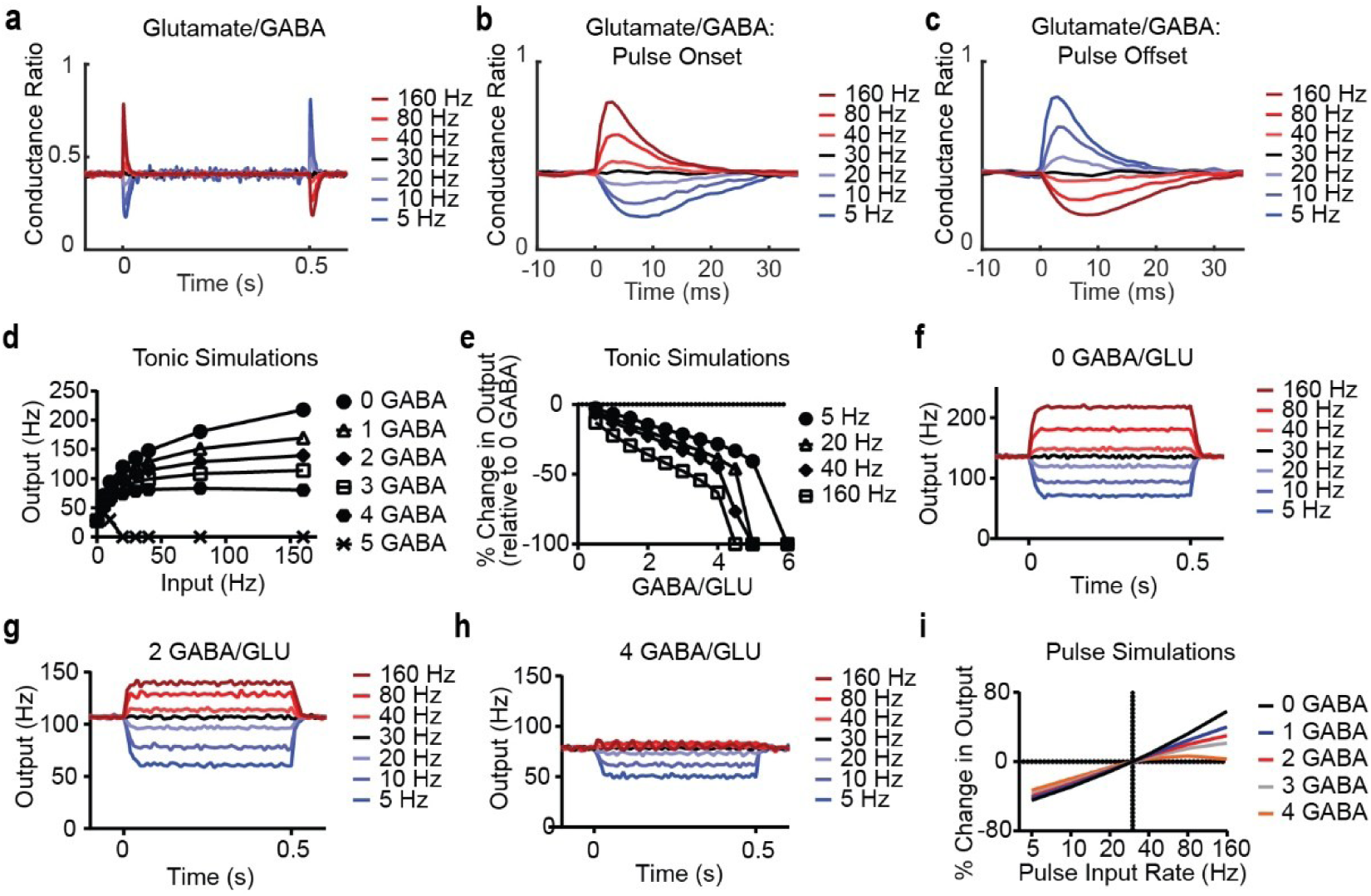
Difference in glutamate and GABA kinetics causes TD-like computations. **a**, Ratio of glutamate-to-GABA conductance during pulse input simulations (average of 1000 simulations), shown for equal peak glutamate and GABA condition. Colors as in Fig. 1g-k,m. **b**, Same as **a**, except showing ratio only during the onset of the pulse input activity. **c**, Same as **b**, but for the offset of pulse input activity. **d**, Input-output curves as in Figure 1e, but during simulations using equal glutamate and GABA kinetics (glutamate kinetics changed to GABA kinetics). **e**, Same data as **d**, but re-calculated and plotted to show the change in output activity for different amounts of GABA co-transmission (relative to no GABA) and different input activity. **f**-**h**, Output activity for different amounts of GABA co-transmission during simulations with artificially equal glutamate and GABA kinetics (average of 1000 simulations). **i**, Same as Figure 1l but during simulations using equal glutamate and GABA kinetics.

### Machine learning classification of synaptic terminals

To determine the prevalence of glutamate/GABA co-releasing terminals in the LHb of mice, rats, and monkeys, we immunolabeled brain slices from each species for markers of glutamate (Vglut2) and GABA (GAD1/2; hereafter, GAD) release that are tightly correlated with glutamatergic and GABAergic transmission in the habenula^2^ (the habenula has little input from Vglut1-expressing terminals, even in monkeys; Fig. S4a-b). Although habenular cell bodies express Vglut2 mRNA^1,60,61^, they contain little Vglut2 and GAD protein (Fig. S4c-d), which are transported to synaptic terminals. After immunolabeling, we made high-resolution, tiled images of the habenula and adjacent thalamus (Fig. 3a-c; N = 36 tiled images from 19 brain slices and 19 mice, 30 tiled images from 22 brain slices and 8 rats, and 23 tiled images from 23 brain slices and 5 monkeys). By eye, although there were similar levels of Vglut2 and GAD co-labeling in the lateral aspect of the LHb among the species (Fig. 3d-f, middle panels), there appeared to be differences in co-labeling in the rest of the LHb (Fig. 3d-f, left panels; Fig. S4c-d), as well as, “ribbon-like” terminal labeling in monkeys but not rodents (Fig. S4c-d).

**Figure 3.**
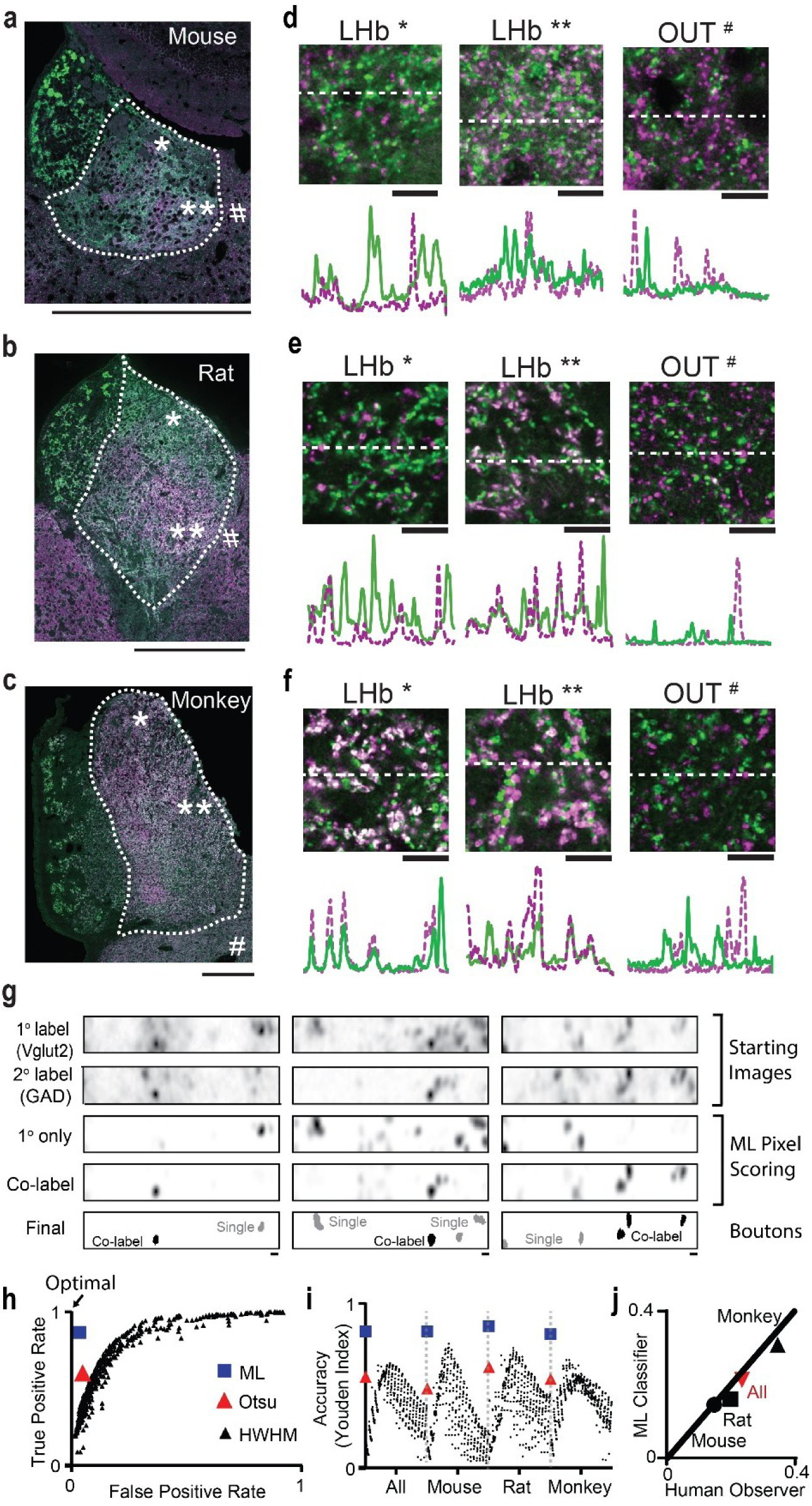
Machine learning classification of glutamate/GABA synaptic terminals. **a**, Example of a high-resolution, tiled image of the mouse habenula. Green, Vglut2. Purple, GAD. Dashed line, LHb. Scale, 0.5 mm. *,**, and # correspond to magnified images in **d**. **b** and **c** are the same as **a**, but for rat and monkey, respectively. **d-f**, Magnified images from the tiled images in **a-c**. Scale, 10 µm; White, co-labeling. Below, line plots of Vglut2 (green) and GAD (purple) signals at the positions indicated by the straight dashed lines. **g**, Examples of machine learning classification of images (left: mouse, middle: rat, right: monkey). The top two rows are images from the same location. “1° only” and “Co-label” images represent scores from the machine learning classifier for each pixel. “Final” images show the final classification of terminals into either single-labeled (gray) or co-labeled (black). Scale, 1 um. **h**, Average ROC plot from images in the training set using three methods of image classification. Multiple thresholds were used for the half-width, half-maximum (HWHM) method and each threshold is shown with a unique data point (see Methods and Fig. S4e). **i,** Youden accuracy scores for each method (symbols as in **h**), shown combined for all species and separately. **j**, Comparison of co-labeling scores for the training set from the human observer and the machine classifier. Circle, Mouse. Square, Rat. Black triangle, Monkey. Red triangle, all combined. Line is x = y.

To quantify co-labeling in synaptic terminals, we initially used fluorescence thresholding methods (Fig. S4e). However, accuracy was inconsistent with these methods (Fig. S4f-g). We therefore trained a machine learning classifier on a small dataset to label pixels as ‘background’, ‘single-labeled’, or ‘double-labeled’, and contiguous non-background pixels were grouped together and classified as single or double-labeled, depending on whether the majority of pixels in the bouton were classified as single or double-labeled (see Methods; Fig. 3g). Classification was done twice for each image - once to quantify the proportion of Vglut2-expressing (Vglut2+) terminals that expressed GAD (GAD+), and a second time to quantify the proportion of GAD+ terminals that were Vglut2+. We then used these proportions to calculate the proportion of all terminals that were co-labeled for GAD and Vglut2 (see Methods). An advantage of this method is that it does not require all terminals in the images to be scored to accurately quantify co-labeling, which would require difficult decisions about which terminals are too faint to include in the analysis.

The performance of the machine learning classifier was better than the other methods (Fig. 3h-i and Fig. S5). Compared to Otsu’s method, a commonly used distribution thresholding algorithm (Fig. S4e), the classifier had both fewer false positives (judging a single-labeled terminal as double-labeled; ML classifier: 3.5%, Otsu: 4.9%; N = 2941, 5652 terminals; Fig. 3h) and fewer false negatives (judging a double-labeled terminal as single-labeled; ML classifier: 13.3%, Otsu: 39.7%; Fig. 3h; using human scoring as “ground truth”). This was true even for images the classifier was not trained on (false positives, ML classifier: 2.6%, Otsu: 4.7%; false negatives, ML classifier: 11.7%, Otsu: 35.9%; N = 873, 1716 terminals). The classifier’s accuracy was also similar for all three species (Fig. 3i and Fig. S5), its co-labeling quantification was similar to human judgment (Fig. 3j), and it could discriminate immediately adjacent Vglut2+ and GAD+ terminals with our optics (Fig. S6).

### Evolutionary rise of glutamate/GABA terminals in the mammalian LHb

We then used the machine learning classifier on the entire dataset to calculate the percentage of synaptic terminals in the LHb and the adjacent thalamus that were co-labeled for Vglut2 and GAD. In all three species, there were more co-labeled terminals in the LHb than the adjacent thalamus, consistent with previous studies in rodents^2,41^ (Fig. 4a; 2-way RM ANOVA: Species x Region, F(2,86) = 54.2, P < .0001; Mouse, LHb vs. Outside, P < .0001, Tukey’s test with multiple comparisons, N = 36 tiled images from 19 brain slices and 19 mice; Rat, LHb vs. Outside, P < .0001, Tukey’s test with multiple comparisons, N = 30 tiled images from 22 brain slices and 8 rats; Monkey, LHb vs. Outside, P < .0001, Tukey’s test with multiple comparisons, N = 23 tiled images from 23 brain slices and 5 monkeys; Terminals: Total, 1,502,099; Mouse, 362,631; Rat, 418,405; Monkey, 721,063). There was also more co-labeling in the rat LHb than the mouse LHb, and more co-labeling in the monkey LHb compared to both the rat and mouse LHb (Fig. 4a; Mouse LHb vs Rat LHb, P < .0001, Tukey’s test with multiple comparisons; Mouse LHb vs Monkey LHb, P < .0001, Tukey’s test with multiple comparisons; Rat LHb vs Monkey LHb, P < .0001, Tukey’s test with multiple comparisons; N same as above), indicating proliferation of glutamate/GABA terminals during evolution. Additionally, there was slightly more co-labeling in the adjacent thalamus of monkeys compared to the adjacent thalamus of rats and mice (Fig 4a; Fig. S7b-c; Mouse Outside vs Monkey Outside, P = .047; Rat Outside vs Monkey Outside, P = .04; Mouse Outside vs Rat Outside, P = 0.98, Tukey’s test with multiple comparisons; N same as above), possibly due to collaterals from basal ganglia inputs that target the LHb - collaterals which are known to exist in monkeys^62^, but not rodents^28^. The species differences in LHb co-labeling remained statistically significant when accounting for possible correlations in scores from the same tissue slices and animals^63^, as well as, potential sex differences (ANOVA (GLME): Species, F = 35.7, P = 5.7 x 10 ^-12^; Sex, F = 1.5, P = 0.2; Mouse vs Rat, t = 2.8, P = .007; Mouse vs Monkey, t = 8.4, P = 7.2 x 10 ^-13^; Rat vs Monkey, t = 5.7, P = 2.1 x 10 ^-7^; Sex, t = 1.2, P = 0.2; N same as above). Although some of the tissue was from monkeys with prior ethanol exposure, this did not explain the higher LHb co-labeling scores in monkeys, because co-labeling tended to be higher in control monkeys without ethanol exposure (GLME: Control: 43.5 %, N = 17 tiled images/slices, 3 monkeys; Prior Ethanol: 29.9%, N = 6 tiled images/slices, 2 monkeys; GLME, t = 2.0, P = .06).

**Figure 4.**
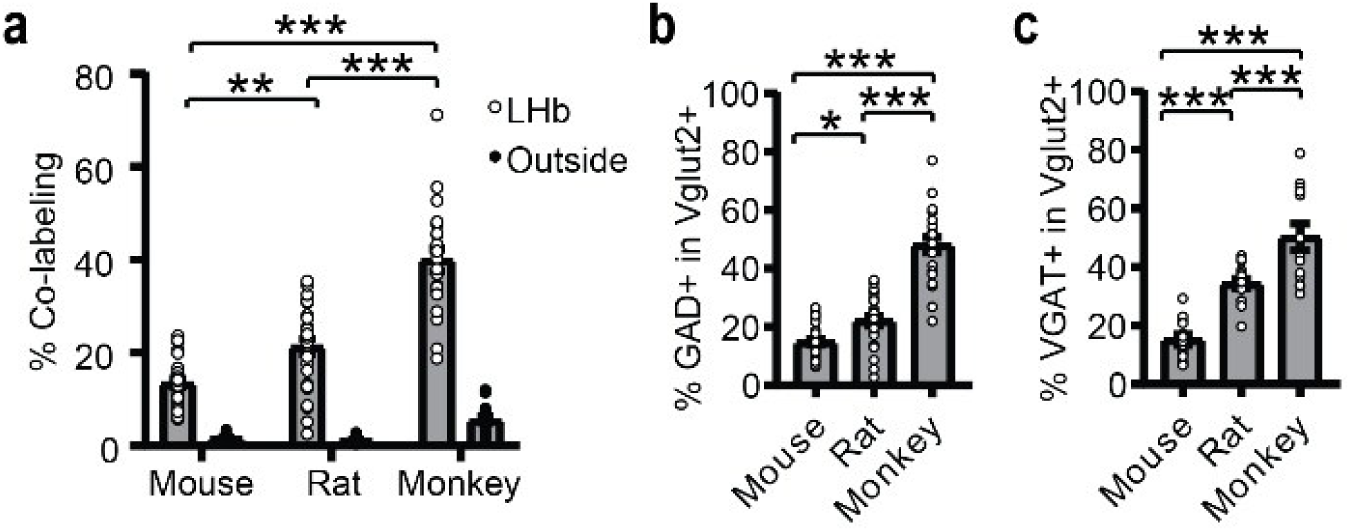
Cross-species quantification of glutamate/GABA synaptic terminals in the habenula. **a**, Co-labeling of Vglut2 and GAD in synaptic terminals from mice, rats, and monkeys. “Outside” refers to the thalamic regions surrounding the LHb. **b**, Percentage of glutamatergic terminals that express GAD in the LHb. **c**, Percentage of glutamatergic terminals that express VGAT in the LHb. *, P < .05; **, P < .01; ***, P < .001.

The species differences in Vglut2/GAD co-labeling were due to differences in the percentage of Vglut2+ terminals that were also GAD+ (Fig. 4b; ANOVA (GLME): Species, F = 47.1, P = 1.6 x 10 ^-14^; GLME: Mouse vs Rat, t = 2.4, P = .02; Mouse vs Monkey, t = 9.6, P = 2.8 x 10 ^-15^; Rat vs Monkey, t = 7.1, P = 3.8 x 10 ^-10^), rather than the percentage of GAD+ terminals that were Vglut2+, which was greater than 50% in all three species (Mouse, 58%; Rat, 81%; Monkey, 69%). Consistent with these results, we found similar species differences in the percentage of glutamatergic terminals that were also GABAergic in a smaller dataset that used the vesicular GABA transporter (VGAT), rather than GAD, as a marker of GABAergic terminals (Fig. 4c and Fig. S7a; ANOVA (GLME): Species, F = 27.5, P = 3.5 x 10 ^-8^; GLME: mouse vs rat, t = 4.3, P = .0001; mouse vs monkey, t = 7.4, P = 6.3 x 10^-9^; rat vs monkey, t = 3.6, P < .001; N = 14 mouse, 15 rat, and 13 monkey tiled images/slices from 6 mice, 8 rats, and 5 monkeys).

To further test for proliferation of glutamate/GABA co-release during evolution, we quantified glutamate/GABA terminals in zebrafish, with the expectation that there would be fewer dual glutamate/GABA terminals in the fish habenula than the mammalian habenula. This result would be consistent with previous studies that reported glutamatergic, but no GABAergic, inputs from the EPN^64–66^ in fish (the EPN is also a major input to the habenula in fish^67^). Consistent with the hypothesis, there was little overlap of Vglut2 and GAD in the fish habenula using the same antibodies we used in mammalian tissue (Fig. S8a-b; Co-labeling: Hb, 2.4 +/- 0.6 %, N = 6 fish; Outside, 1.5 +/- 0.7 %, N = 7 fish, t test, P = .34; vHb, 3.1 % +/- 1.0%, N = 6 fish; dHb, 1.7 +/- 0.7 %, N = 6 fish; paired t test for vHb vs dHb, P = 0.52; paired t test for vHb vs Outside, P = 0.19), and less co-labeling than in the mouse LHb (Fig. S8c; t test, P < .0001; fish vHb vs mouse LHb, t test, P < .0001; N = 6 fish and 19 mice). However, we did observe a small, but statistically significant, increase in the percentage of Vglut2+ terminals that were also GAD+ in the fish habenula compared to adjacent regions when combining terminals from all fish together (Fig. S8d; chi-square = 3.9, P = .049; N = 354 habenula and 1051 “Outside” terminals).

### Glutamatergic EPN neurons in fish express fewer GABAergic markers than glutamatergic EPN neurons in mice

To complement the fish immunohistochemical results, we analyzed published single-cell RNA-sequencing data from the zebrafish EPN^64^ and compared them to those from the mouse EPN^28^. Like mammals^28^, fish have a glutamatergic EPN population that projects to the habenula and a GABAergic population that projects outside the habenula, including the thalamus^64–68^. Compared to non-glutamatergic (i.e., Vglut2-negative) EPN neurons in fish, glutamatergic (i.e., Vglut2+; there were no Vglut1+ cells in the EPN) EPN neurons in fish were less likely to express GABAergic markers (Fig. 5a; Fisher’s tests for GAD1, GAD2, and VGAT, all P < .0001), consistent with primarily glutamatergic transmission from this population ^65–67^. In contrast, Vglut2+ EPN neurons in mice were equally likely to express GABAergic markers as Vglut2-negative EPN neurons (Fig. 5b; Fisher’s tests for GAD1, GAD2, and VGAT, all P > .05), consistent with an increase in glutamate/GABA EPN neurons in mammals. To directly compare fish and mouse glutamatergic EPN populations, we controlled for differences in expression counts across the genome by downsampling the expression counts of all genes per cell in the mouse dataset to match the median counts per cell in the fish dataset (see Methods). Despite downsampling the mouse gene expression counts, we found that Vglut2+ EPN neurons in fish were much less likely to express GABAergic markers than Vglut2+ EPN neurons in mice (Fig. 5c; Fisher’s tests for GAD1, GAD2, and VGAT, all P < .0001), and Vglut2+ EPN neurons in fish had less mean expression of GABAergic markers, but not Vglut2 (Fig. 5d; Wilcoxon rank sum tests for GAD1, GAD2, and VGAT, all P < .0001, for Vglut2, P = .62). These data further support proliferation of glutamate/GABA co-release in LHb circuits during evolution.

**Fig. 5.**
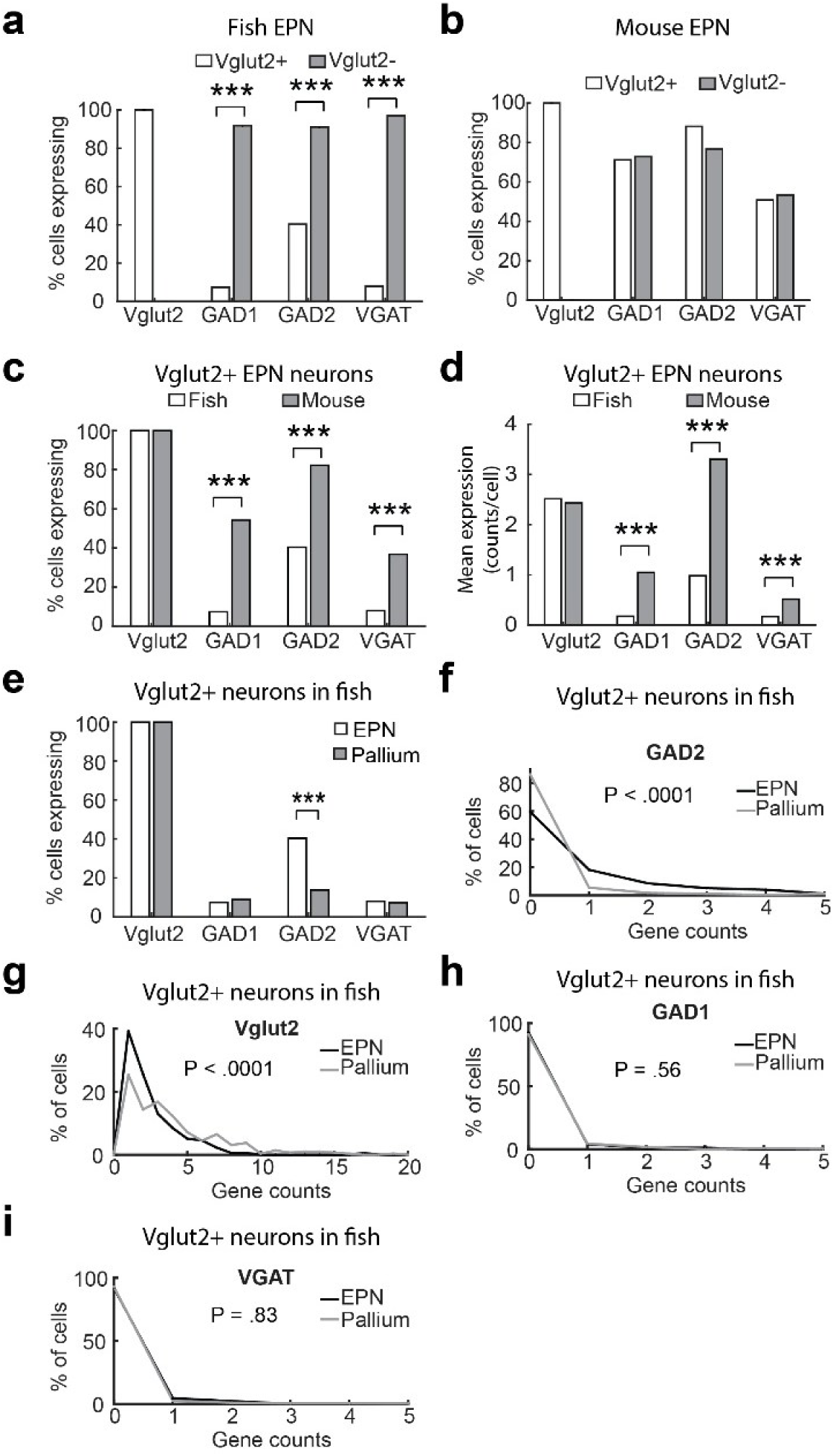
Glutamatergic EPN neurons in fish express fewer GABAergic markers than those in mice. Analysis of single-cell RNA-sequencing data from Wallace et al., (2017) and Tanimoto et al., (2024). **a**, Comparison of gene expression from zebrafish EPN Vglut2+ cells (N = 176) and EPN Vglut2- cells (N = 1477). **b**, Same as **a**, but for mouse (N = 59 EPN Vglut2+ and 107 EPN Vglut2- cells). **c**-**d**, Comparison of gene expression from zebrafish (N = 176) and mouse (N = 51) Vglut2+ EPN neurons, as indicated. Note that mouse gene expression for each cell was downsampled across the genome to match the median zebrafish genome expression for each cell. **e**-**i**, Comparison of gene expression from zebrafish EPN Vglut2+ cells (N = 176) and pallial Vglut2+ cells (N = 292). *** P < .0001, Fisher’s test (% cells expressing comparisons) and Wilcoxon rank sum test (mean expression).

To test whether the zebrafish EPN has any glutamate/GABA co-expression phenotype, we compared GABAergic marker expression in EPN Vglut2+ neurons to that in pallial (cortex-like) Vglut2+ neurons in zebrafish from the same dataset. We found that EPN Vglut2+ and pallial Vglut2+ neurons were equally unlikely to express GAD1 and VGAT (Fig. 5e,h,i; Fisher’s and Wilcoxon tests, P > .05); however, EPN Vglut2+ neurons were more likely to express GAD2 than pallial Vglut2+ neurons (Fig. 5e,f; Fisher’s and Wilcoxon tests, P < .0001) suggesting the existence of a small population of glutamatergic EPN neurons with partial GABAergic identity in fish.

### Altered topography of glutamate/GABA terminals in the primate LHb

The large number of analyzed terminals allowed us to map the topography of glutamate/GABA terminals in the LHb of each mammalian species (see Methods; Fig. 6a-c). In both mice and rats, there was a dominant medial-lateral gradient, consisting of more GAD co-labeling in glutamatergic terminals in the lateral LHb compared to the medial LHb, as expected from the known topography of glutamate/GABA co-releasing EPN inputs to the LHb^28,41^ (Fig. 6d-e, n, Fig. S9d; GLME: Mouse, lateral LHb vs. medial LHb, t = 7.7, P = 6.6 x 10 ^-11^, N = 35 tiled images from 19 slices from 19 mice; Rat, lateral LHb vs. medial LHb, t = 12.9, P = 1.1 x 10 ^-18^, N = 30 tiled images from 22 slices from 8 rats). Surprisingly, there was not more co-labeling in the lateral LHb than the medial LHb in monkeys (Fig. 6f, g, Fig. S9d; co-labeling was shifted slightly medially, GLME: t = 2.6, P = .013, N = 22 tiled images from 22 slices from 5 monkeys). Instead, there was a dominant ventral-dorsal gradient in monkeys, with more co-labeling in the evolutionarily newer^69^, dorsal LHb than the ventral LHb (Fig. 6f, h, Fig. S9d; GLME: t = 5.7, P = 9.6 x 10 ^-7^, N = 22 tiled images from 22 slices from 5 monkeys), consistent with expansion of glutamate/GABA co-release during evolution. These topographical differences were replicated in the smaller VGAT dataset (Fig. S9a-c; GLME: Mouse, lateral LHb vs. medial LHb, t = 7.8, P = 3.0 x 10 ^-8^, N = 14 tiled images/slices from 6 mice; Rat, lateral LHb vs. medial LHb, t = 15.4, P = 1.4 x 10 ^-14^, N = 14 tiled images/slices from 8 rats; Monkey, dorsal LHb vs ventral LHb, t = 5.8, P = 5.8 x 10 ^-6^, N = 13 tiled images/slices from 5 monkeys; other within-species comparisons, P > .05).

**Figure 6.**
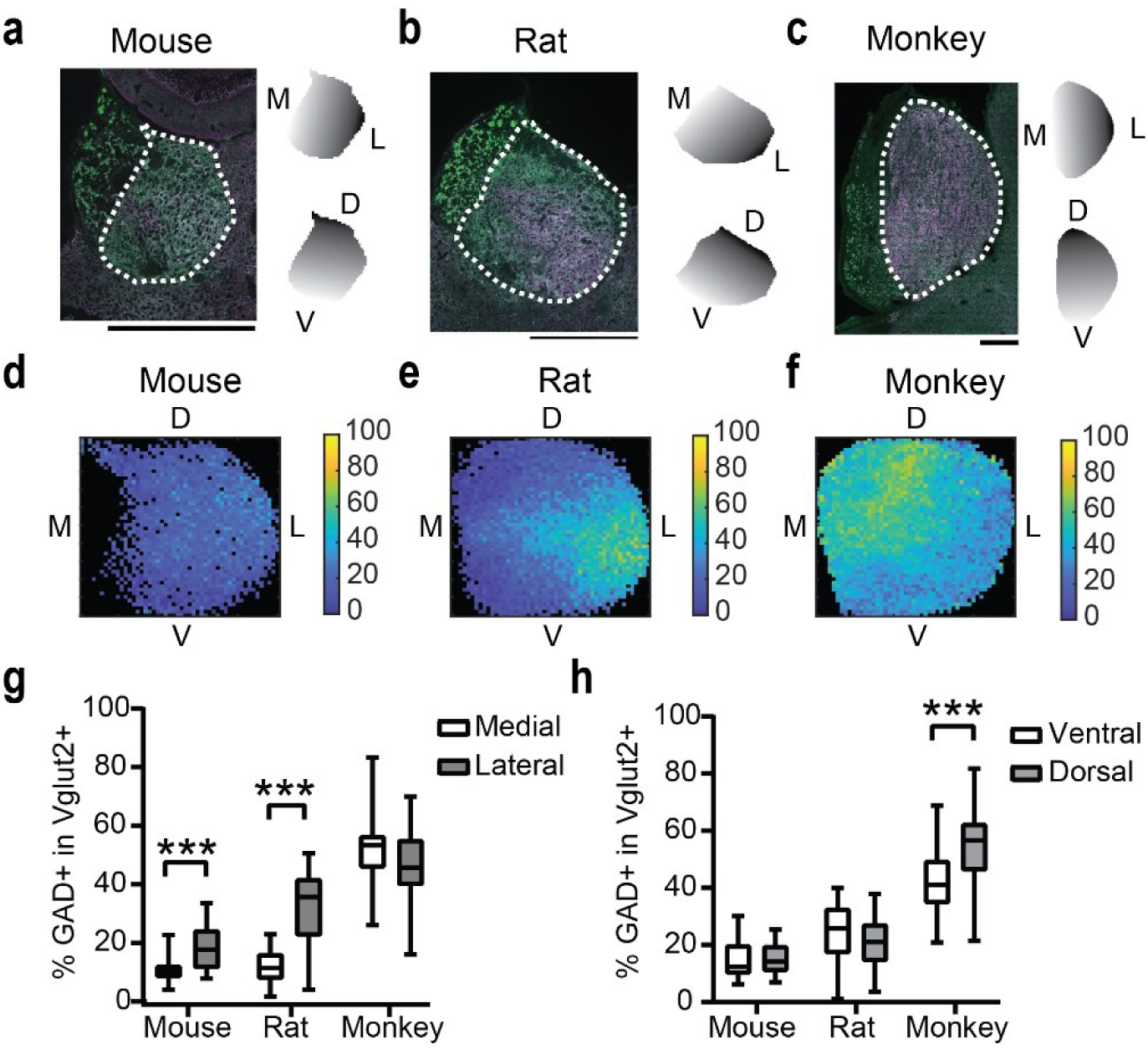
Expansion of glutamate/GABA terminal topography in the primate LHb. **a**, Left, Example high-resolution tiled image of the mouse habenula, as in Figure 3. Right, Shading shows medial-lateral and dorsal-ventral gradients used to quantify the topography of co-labeling. **b** and **c** are the same as **a**, but for rat and monkey, respectively. **d**, Topographical map of the percentage of glutamatergic terminals that express GAD in the mouse LHb (average of 35 tiled images; tiles with data from < 6 tiled images are black). **e**, same as **d** but for rats (average of 30 tiled images). **f**, same as **d** but for monkeys (average of 22 tiled images). **g**, Percentage of glutamatergic terminals that express GAD in the medial LHb and lateral LHb for each species (same tiled images as in **d**-**f**). **h**, same as **g**, but for the dorsal and ventral LHb. ***, P < .001.

## Discussion

Here we show that GABA co-release with glutamate is sufficient to produce TD-like transformations of input activity (i.e., differentiation; Fig. S10) due to the slower kinetics of GABAergic transmission. Our data also indicate that glutamate/GABA terminals evolutionarily proliferated from fish to monkeys and expanded topographically in the primate LHb - a subcortical region that computes TD-like reward prediction errors to drive learning^13,21,44,49,70–74^. Therefore, GABA co-release with glutamate may create TD-like reward prediction errors in the LHb – a region which may be uniquely positioned to exploit this synaptic mechanism due to its high basal firing rate, connectivity, and apparent lack of GABAergic interneurons^61,75,76^ (but see ^77,78^).

TD reinforcement learning theory predicts that differentiation should act on value signals, and studies indicate that basal ganglia inputs to the LHb^23,24^ instead already convey TD error-like activity (some EPN co-releasing inputs may encode action value^27^, but this remains to be tested directly). Thus, it is possible that the primary function of GABA co-release from the EPN to the LHb is to diversify TD error-like activity, rather than create it. Future studies, including studies which measure TD error-like signals during perturbation of GABA co-transmission, are needed to determine if inputs to the LHb, including the VTA or basal forebrain^4,5,8^, use GABA co-transmission to create TD error-like signals in the LHb. It is also important to note that although the LHb is a hub for this unusual form of synaptic transmission in mammals^1,4^, miniature post-synaptic current analysis suggests that GABA co-release with glutamate is more widespread in the mammalian brain than previously thought^79^. Therefore, it is possible that GABA co-release with glutamate is used for differentiation outside of the LHb, and even upstream of basal ganglia inputs to the LHb.

Recent studies discovered that VTA dopamine neurons have heterogeneous TD error-like signals that are predicted to contribute to distributional learning and complex decision-making^31,34,35^. Habenula lesions disrupt TD error-like signals in VTA dopamine neurons, alter the basal firing rate of VTA dopamine neurons, and increase anticipatory licking and relative VTA dopamine neuron responsiveness to cues that probabilistically predict reward^22,32^. Our simulations indicate that GABA co-release may skew the activity of LHb neurons (and therefore also dopamine neurons) to produce asymmetric responses to negative (increases in input activity) and positive feedback (decreases in input activity). This is consistent with a prior study that found an important role for the LHb in flexible, higher-order decision-making and other studies showing that a line of rats with reduced GABA co-release from the EPN to the LHb have altered preference for a “risky” response option^2,20,80^. It is also consistent with habenula-VTA functional coupling affecting risk preferences during learning in humans^81^. Together, these data suggest that glutamate/GABA co-transmission in the LHb contributes to heterogeneous TD error signaling in VTA dopamine neurons and higher-order decision-making, perhaps through topographic projections to inhibitory neurons in the brainstem^13–20^.

Multiple lines of evidence indicated changes of glutamate/GABA circuits into the LHb during evolution, including Vglut2/GAD and Vglut2/VGAT terminal co-labeling in multiple species, altered and expanded topography of Vglut2/GAD and Vglut2/VGAT terminal co-labeling in monkeys towards the evolutionarily newer dorsal LHb^69^, increased variation in the ratio of glutamate/GABA conductance in rats compared to mice, and increased GABAergic marker expression in mouse relative to fish Vglut2+ EPN neurons, a major input to the habenula in fish and mammals^67,82^. The single-cell RNA-sequencing analysis further suggested the existence of a small Vglut2+ EPN population with a partial GABAergic identity in fish, because fish EPN Vglut2+ neurons were more likely to express GAD2, but not VGAT nor GAD1, than pallial Vglut2+ neurons from the same dataset. These data suggest that this population may be a precursor to the mammalian glutamate/GABA EPN population.

Proliferation of glutamate/GABA terminals during mammalian evolution is also consistent with a role for glutamate/GABA co-transmission in flexible, higher-order learning and decision-making. However, the precise significance of this increased abundance of glutamate/GABA terminals is unclear. An increased use of glutamate/GABA co-transmission may allow greater diversity of TD error-like signals^31,35,83^ and therefore a richer representation of value in downstream circuits^34^. Consistent with increasing diversity among LHb neurons, we found larger variation in glutamate/GABA co-transmission, concomitant with more glutamate/GABA LHb terminals, in rats than mice.

More glutamate/GABA terminals in the LHb also may reflect greater dedication of mammalian brains to learned behavior that depends on dopaminergic reinforcement signals. Consistent with this possibility, fish lack a midbrain dopaminergic system^84^ (but see ^85,86^) and dopaminergic input to the cortex is denser in primates than rodents^87^. It will be important to determine which LHb inputs are responsible for the expansion of glutamate/GABA terminals to the dorsal, evolutionarily newer^69^ part of the LHb in primates, as well as, the projection targets of neurons in this region of the primate LHb and whether they are more likely to encode TD error-like activity than neurons in other parts of the LHb. This information may yield clues about the evolution of LHb function, a subcortical brain region which is often regarded as highly conserved in vertebrates.

## Methods

### Animals

Four adult male rhesus macaques, 7.5 years old at time of sacrifice, and one female cynomolgus macaque, 8.5 years old at time of sacrifice were used (all obtained from MATRR). Two of the four male macaques had prior ethanol consumption, which ended at least 56 days prior to sacrifice. Nineteen adult C57Bl/6J mice (9 female; 3-6 months old; 1-4/cage; 12/12 hour light-dark cycle) were used for Vglut2/GAD immunohistochemistry. Eight of 19 mice used for Vglut2/GAD immunohistochemistry had prior footshock, but were grouped together with the non-shocked mice because we found no differences in co-labeling between the shocked and non-shocked mice. Eight adult C57Bl/6N mice (4 female; 1-4/cage; 12/12 hour light-dark cycle) were used for Vglut2/VGAT immunohistochemistry. Eight adult Sprague-Dawley rats (4 female; 2-5 months old; 1-4/cage; 12/12 hour light-dark cycle) were used for Vglut2/GAD and Vglut2/VGAT immunohistochemistry. Seven adult zebrafish (WIK strain; 6 months old; mix of male and female) were used for Vglut2/GAD immunohistochemistry. For other ex vivo electrophysiology experiments, we used heterozygous, adult Sst-cre mice (JAX:028864; 1- 4/cage), C57Bl/6 mice (1-4/cage), and Sprague-Dawley rats (1-4/cage). All procedures involving fish and rodents were approved by the Institute Animal Care and Use Committee of the University of Texas Southwestern Medical Center.

### Immunohistochemistry

Mice and rats were anesthetized with an overdose of ketamine/dexmedetomidine and transcardially perfused with 0.9% NaCl followed by 4% formaldehyde (10% formalin) solution. Brains were post-fixed for 1-2 days and then stored in 30% sucrose until slicing. Monkeys were anesthetized with ketamine (15 mg/kg, i.m.) and sodium pentobarbital (30-50 mg/kg, i.v.) and sacrificed and perfused as described in ^88^ (cold ACSF, then -80 C storage). Frozen chunks of monkey tissue with the habenula were submerged in 4% formaldehyde for 1-2 days, then increasing concentrations of sucrose (10, 20, 30%) and stored at 4 °C until preparation for slicing. Fish were put in ice water for 10 minutes, then decapitated and stored in 4% paraformaldehyde for 1-2 days, followed by PBS at 4 °C. Mouse, rat, and monkey tissue were sliced coronally at 30 um on a cryostat. Fish brains were embedded in low-melting point agarose and sliced coronally at 50-75 um on a vibratome. Slices were washed in PBS, then put in 1X sodium citrate antigen retrieval solution at 80 °C for 30 minutes, then immunolabeled with primary antibodies (GAD1/2, ThermoFisher PA1-84572: 1:1000 or VGAT, Millipore AB5062P: 1:500 and Vglut2, Millipore MAB5504: 1:500; Vglut1, Novus, NBP2-46627: 1:2000, 0.2% Triton, 3% normal goat serum) for 2 days at 4 °C, then secondary immunolabeling for 3 hours at room temperature (1:250 goat anti-rabbit Alexa 568 and goat-anti-mouse 488 or goat-anti-mouse 647), followed by washing and mounting with Vectashield Anti-fade with DAPI). Controls without primary antibodies showed no labeling.

### Imaging

High-resolution (1024×1024), tiled images of the habenula were taken with a 63x objective and Zeiss LSM 800 confocal microscope. Laser power and PMT gain were set to minimize background fluorescence and avoid saturation. Pinhole size was set to 1 airy unit (using the red channel) and tile overlap was 1% for image stitching. TIFF files were exported for MATLAB analysis. All tiled images in the manuscript are arranged with medial on the left and dorsal on top.

### Surgery

For ex vivo electrophysiology, mice were anesthetized with isoflurane for stereotaxic bilateral injection of 0.2-0.3 µL AAV virus into the EPN (AAV2/8-ChR2(H134)-EYFP or AAV2/5- DIO-ChR2(H134)-EYFP for Sst-cre mice; A-P: -1.15 mm from bregma; M-L: 1.8 mm; D-V: -4.3 - -4.5 mm from skull surface at bregma). To minimize pain after surgery, all animals were subcutaneously injected with analgesic carprofen (Rimadyl, 1.3mg/mL, 0.1mL per 25g). Location of the injections was confirmed by visualization of the fluorophore expressed by the virus.

### Image analysis

1024×1024 TIFF files were imported into MATLAB, smoothed using the imgaussfilt function (sigma = 2), and arranged in two dimensions to match the original tiled image. An outline of the lateral habenula (distinguished from the medial habenula by cell density and tissue morphology) and regions outside the habenula, as well as a dorsal-ventral orientation line (parallel to the longitudinal orientation of the medial habenula), were made using custom MATLAB code. Each tile from mouse, rat, and fish (1024 x 1024) was divided into 64 128 x 128 sub-tiles. Monkey Vglut2/GAD tiles were divided into 16 256 x 256 sub-tiles for further analysis and monkey Vglut2/VGAT tiles were divided into 64 128 x 128 sub-tiles. For the topography maps, the location of each sub-tile was normalized to its location in the dorsal-ventral and medial-lateral axes (-1 to 1, and subsequently, 0 to 1) using custom code. A subset of 30 x 256 images was selected randomly from the mammalian tissue set for testing different methods of quantifying overlap of Vglut2 and GAD: half-width, half-max (HWHM), Otsu’s method, and machine learning (ML). The HWHM method sets thresholds for each label according to the distance from the peak of the signal distribution, in HWHM units (see Fig. S4e). The pixel intensity distribution for each training image was derived from the entire tiled image that the training image was drawn from. For each immunolabel (e.g., Vglut2 or GAD), 20 HWHM-derived thresholds were set at 0.25 HWHM intervals - three below the peak, one at the peak, and sixteen above the peak, thus covering 0.75 HWHM units below the peak and 4.0 HWHM units above the peak of the intensity distributions. Because there were 20 HWHM thresholds for each of two immunolabel signals, there were a total of 400 (20*20) combinations of HWHM thresholds tested. Otsu’s method derives a single threshold for each immunolabel signal by minimizing the intraclass variance after splitting the tiled image pixel intensity distribution in two.

Our machine learning classifier was trained using stochastic gradient descent in MATLAB (using trainNetwork) on 70% of the image subset, using a 0.1 dropout rate and .001 learning rate that reduces by 5% across training epochs. Initially, multiple classifiers were trained and the one with the best accuracy was selected for continued training. The classifier categorizes each pixel of a 32×256 image as background, single-labeled (only the primary label, e.g., Vglut2), or double-labeled (both the primary and secondary label, e.g, both Vglut2 and GAD). To do so, it is fed 32×256×4 images consisting of the original primary label 32×256 image (smoothed using a Gaussian filter), the original, smoothed secondary label 32×256 image of the same region, a re- scaled, smoothed, primary label 32×256 image (from 0 to 1, with 1 equal to maximum intensity of that label for the 32×256 image), and a re-scaled (0 to 1), smoothed, secondary label 32×256 image. Contiguous, non-background pixels were then grouped together into boutons and classified as co-labeled if the majority of pixels in the bouton were classified as co-labeled by the machine learning classifier.

The training image set was made by random selection of 30×256 regions from the mammalian images. Each image was then analyzed by a human observer using a custom MATLAB program that kept the observer blind to the type of signal and origin of the image. The program would display a 30×256 image for one label only (primary label) and ask the observer to circle clear synaptic terminals. Then this image would disappear, and the corresponding 30×256 image for the secondary label would appear with circles where the observer had drawn them for the primary label. The observer would then enter their judgment about whether there was a secondary- labeled synaptic terminal in each circle or not (if the observer was unsure, the circled region would be ignored).

Scoring accuracy via HWHM, Otsu, and machine learning methods was quantified by calculating the true positive rate (TPR; proportion of double-labeled terminals, as judged by human observation, that were judged to be double-labeled) and the false positive rate (FPR; proportion of single-labeled terminals that were judged to be double-labeled). The Youden index (TPR - FPR) was used as a measure of accuracy for each measure.

Co-labeling percentage was calculated using:

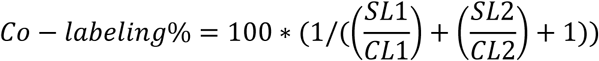

where SL1/CL1 is the ratio of single labeled terminals for immunolabel #1 as the primary label to co-labeled terminals using immunolabel #1 as the primary label, and SL2/CL2 is the ratio of single labeled terminals for immunolabel #2 as the primary label to co-labeled terminals using immunolabel #2 as the primary label.

All slices were included in all analyses except one mouse slice (Vglut2/GAD dataset), one monkey slice (Vglut2/GAD dataset), and one rat slice (Vglut2/VGAT dataset) that were excluded from the topography analyses because of either tissue distortion/tears (mouse and rat slices) or D-V orientation uncertainty (monkey slice), as well as, one monkey slice in the Vglut2/VGAT dataset that was excluded from all analyses because of autofluorescence.

### Neural simulations

Simulations were performed in NEURON using custom code written in Python and were designed to model GABA co-release with glutamate from the major input to the LHb from the basal ganglia, for which there are data on single-cell firing rates and single-cell responses to rewarding and aversive stimuli. We used a minimal circuit model to determine how GABA co-release with glutamate changes neuronal output, consisting of one output neuron with five dendrites and ten or twenty basal ganglia input neurons, as indicated. We used NetStim units as input neurons with noise = 1 (maximal randomness), meaning presynaptic interspike intervals are randomly chosen from an exponential distribution with mean equal to the inverse of the mean firing rate, thus simulating a Poisson process. Unless stated otherwise, the output neuron contained only voltage-gated sodium and potassium channels, leak conductance, AMPA receptors, and GABA-A receptors. Each input neuron synapsed onto a different part of the output neuron’s dendritic tree, with synapses spaced 25 um (20 input simulations) or 50 um (10 input simulations) apart. The number of independent basal ganglia axons that connect to each LHb neuron is unknown and was therefore estimated from spontaneous, action-potential mediated, mixed glutamate/GABA conductances and the maximal, bulk, conductance from synchronized basal ganglia axon stimulation. Single input AMPA-R mediated peak conductance was .004 uS in all simulations, which was estimated from spontaneous, action-potential mediated, mixed glutamate/GABA conductances. In all simulations, to mimic the high basal firing rate and depolarized resting potential of LHb neurons, constant current was injected into the output neuron to increase its basal firing rate to ∼ 30 Hz without input stimulation. AMPA-R and GABA-A-R conductance kinetics parameters were chosen to match the average experimentally observed AMPA-R and GABA-A-R kinetics at physiological temperature. We used the same GABA-B modeling and kinetics as in ^89^, with conductance amplitude set to 10% of AMPA-R and GABA-A-R conductances. For calculating % change in output during pulse simulations, output activity in the first 200 ms after the start of the pulse was compared to the pre-pulse period.

### Electrophysiology

Two to five weeks after surgery, mice and rats were anesthetized with isoflurane before decapitation and brain removal. Brains were chilled in ice-cold dissection buffer (110.0 mM choline chloride, 25.0 mM NaHCO_3_, 1.25 mM NaH_2_PO_4_, 2.5 mM KCl, 0.5 mM CaCl_2_, 7.0 mM MgCl_2_, 25.0 mM glucose, 11.6 mM ascorbic acid, 3.1 mM pyruvic acid; gassed with 95%O_2_/5%CO_2_) and cut in 300-400 micron thick coronal slices through the entopeduncular nucleus and LHb. Slices were transferred to 35°C ACSF (118 mM NaCl, 2.5 mM KCl, 26.2 mM NaHCO_3_, 1 mM NaH_2_PO_4_, 20 mM Glucose, 1 mM MgCl_2_, 2 mM CaCl_2_; 22°–25°C; pH 7.4; gassed with 95%O_2_/5%CO_2_) for 30 minutes. After an additional 30 minutes of recovery at room temperature, slices were transferred to the recording chamber and constantly perfused with room temperature (22-25 °C) ACSF. Some of these whole-cell recording data from mice and all of the whole-cell recording data from rats were previously used in ^2^.

For experiments in Sst-cre mice (sacrificed 3-5 weeks after surgery), mice were injected with ketamine and dexmedetomidine and rapidly perfused with 10-15 mL cold dissection buffer, as above, that included 2 mM kynurenic acid. Brains were removed and cut in cold dissection buffer and transferred to 35°C ACSF with 1 mM ascorbic acid for 30 minutes. After an additional 30 minutes of recovery at room temperature, slices were transferred to the recording chamber and constantly perfused with physiological temperature (37 °C) ACSF.

Recordings were made from cells in the lateral half of the rodent LHb, where entopeduncular inputs were densest. The intracellular solution consisted of (in mM): 7.5 QX314, 115 cesium methanesulfonate, 20 CsCl, 10 HEPES, 2.5 MgCl_2_, 4 Na_2_-ATP, 0.4 Na-GTP, 10 Na-phosphocreatine, and 0.6 EGTA (pH 7.2). 470 nm light pulses were delivered to the LHb via an LED. Isolated AMPA-R-mediated or GABA-A-R-mediated synaptic responses were obtained in the presence of D-APV (100 µM) in voltage-clamp mode near the reversal potentials of GABA-A-R and AMPA-R currents, respectively. For data in Figure 1a and Fig. S1a, neurons with small optogenetic responses were excluded from all analyses (if both AMPA-R and GABA-A-R peak currents were < 100 pA).

### Single-cell RNA-sequencing analysis

Single-cell gene expression data from the zebrafish (clusters vEN and EN1-6; left and right hemispheres from 6 fish) and mouse (clusters 5 and 6) EPN were downloaded from ^64^ and ^28^. The fish dataset was in raw gene counts but the mouse dataset needed to be converted back from natural log values to raw gene counts. Fish Vglut2a and Vglut2b isoforms were added together into one Vglut2 category, and Gad1a and Gad1b isoforms were added together into a GAD1 category. For direct comparison of fish and mouse Vglut2+ EPN neurons, we downsampled randomly across the mouse genome for each neuron so that mouse neurons with higher total expression levels than the median from fish Vglut2+ EPN neurons now had the same total number of gene counts as the median gene count from fish Vglut2+ EPN neurons. This random downsampling procedure was repeated 1000 times and the average gene counts for each mouse EPN neuron across the 1000 iterations were used to compare to the fish dataset. Please note that without this downsampling procedure the difference between GABAergic marker expression in Vglut2+ EPN neurons in mice and fish would have been even larger. For the comparison between Vglut2+ EPN neurons in the fish EPN and Vglut2+ neurons in the fish pallium, we also downloaded Pallial1-Pallial4 clusters from ^64^.

### Confocal resolution analysis

Lateral and axial resolution of our optics and images were determined empirically using 50 nm green fluorescent beads (Cat# 17149-10; Polysciences) using the same 63x objective and settings as was used for our tissue imaging. For lateral resolution FWHM calculations, isolated beads were detected as regional intensity maxima, then analyzed from raw 11 × 11 pixel bead-centered regions of interest. Each bead was fit with a two-dimensional Gaussian with independent x and y widths and a constant background offset. Lateral FWHM was calculated separately for x and y as 2.35 × σ × pixel size (where σ is the fitted Gaussian standard deviations), and the bead’s lateral FWHM was defined as the mean of the x and y FWHM values. For axial resolution, isolated beads were analyzed across Z stacks acquired at 100 nm spacing. Axial fluorescence profiles were fit with one-dimensional Gaussian functions, and axial FWHM values were calculated in the same manner. The AF568 point-spread function was modeled as 1.16-fold broader in both dimensions, based on the relative emission wavelength of AF568 to AF488.

### 3D adjacent bouton simulations and analysis

To estimate whether spatially adjacent single-labeled boutons could be erroneously classified as co-labeled, we generated synthetic 3D bouton pairs and analyzed them with the same pipeline used for experimental images. Boutons were modeled as fluorescent spheres with diameters ranging from 0.8 to 1.6 µm in 0.2 µm increments. Each simulated pair contained one AF488-labeled sphere and one AF568-labeled sphere. The AF488 sphere was placed in the focal plane, whereas the AF568 sphere was placed at defined 3D edge-to-edge distances from the AF488 sphere. Distances ranged from 0 to 0.5 µm in 0.1 µm increments, where 0 µm indicated that the two spheres were touching in 3D. The center-to-center distance between spheres was therefore equal to the bouton diameter plus the specified edge-to-edge distance. We also included a 100% overlapping positive control condition, where the AF488 and AF568 spheres were completely overlapping in space.

For each condition, the AF568 sphere was placed in a random 3D direction relative to the AF488 sphere, producing a mixture of lateral and axial offsets. The AF488 sphere was kept at z = 0, and the AF568 sphere could be displaced laterally and/or axially depending on the sampled 3D direction. Four bouton pairs were placed into each simulated 32 × 256 pixel image while preventing overlap between different simulated pairs in the 2D projection. Ten random images were generated for each combination of bouton diameter and 3D separation. AF488 and AF568 intensity were set to an intensity of 50 arbitrary units.

Fluorescence from each sphere was determined by numerically integrating through the sphere in 25 nm axial steps. For each axial slice, the circular cross-section of the sphere was projected into the image plane and weighted by the fluorophore-specific axial point-spread function. The resulting projection was then blurred laterally with a Gaussian point-spread function. The AF488 point-spread function had lateral and axial FWHM values of 0.27 µm and 0.72 µm, respectively. The AF568 point-spread function was modeled as 1.16-fold broader in both dimensions, based on the relative emission wavelength of AF568 to AF488, giving lateral and axial FWHM values of 0.315 µm and 0.835 µm, respectively. Each sphere was normalized so that an in-focus sphere had the specified peak intensity of 50 units.

To make the simulations more realistic, the synthetic AF488 and AF568 bouton signals were added to real background fluorescence images from each channel. The resulting simulated two-channel images were then processed exactly as experimental images. Because the true location and identity of each simulated bouton was known, false co-labeling was quantified specifically within the projected pixels of the AF488-only bouton. For each simulated AF488 bouton, the fraction of its pixels classified by the network as “both” was calculated. An AF488 bouton was counted as falsely co-labeled if at least 50% of its pixels were classified as “both,” matching the criterion used for bouton-level classification in experimental images.

### Statistics

Except where indicated, we used a generalized linear mixed-effects model (GLME), as recommended by ^63^, in MATLAB. This method accounts for possible correlations among data coming from the same slice and animal by modeling them as random effects and is thus usually more conservative than other statistical tests, which assume statistical independence among data from the same slice/animal. In all cases reported here, statistical significance from the GLME was replicated with standard ANOVA and post-hoc tests. All relevant statistical tests are two-tailed. Error bars represent standard error of the mean in all figures, except box and whisker plots, where the error bars show the entire range, the box shows the interquartile range, and the horizontal line shows the median.

## Acknowledgments

We thank Chunfeng Tan and Carol Tamminga for help with immunohistochemistry, Robert McDougal, Ted Carnevale, Joao Moreira, and Salvador Dura for help with NEURON simulations, Vanessa Jimenez, Nikki Walter, Jim Daunais and MATRR for supply and help with monkey tissue, Kevin Dean and Tai Ngo for fluorescent beads, and Peter Dayan and Robert Malinow for valuable comments on a previous version of the manuscript. Research reported here was supported by the Whitehall Foundation, UT Southwestern Medical Center, and the NIH (R01MH120131).

## Author Contributions

N.R.S., L.R., V.R., A.H., I.C., and S.S. collected data. N.R.S., Y. L., Y.S., N.W., R.R., V.N., and S.S. analyzed data. N.R.S. and S.S. conceived the neural simulations. S.S. performed the neural simulations and wrote the manuscript.

## References

1. Yao, Z., van Velthoven, C.T.J., Kunst, M., Zhang, M., McMillen, D., Lee, C., Jung, W., Goldy, J., Abdelhak, A., Aitken, M., et al. (2023). A high-resolution transcriptomic and spatial atlas of cell types in the whole mouse brain. Nature 624, 317–332. 10.1038/s41586-023-06812-z.

2. Shabel, S.J., Proulx, C.D., Piriz, J., and Malinow, R. (2014). Mood regulation. GABA/glutamate co-release controls habenula output and is modified by antidepressant treatment. Science 345, 1494–1498. 10.1126/science.1250469.

3. Root, D.H., Zhang, S., Barker, D.J., Miranda-Barrientos, J., Liu, B., Wang, H.L., and Morales, M. (2018). Selective Brain Distribution and Distinctive Synaptic Architecture of Dual Glutamatergic-GABAergic Neurons. Cell Rep 23, 3465–3479. 10.1016/j.celrep.2018.05.063.

4. Xu, J., Jo, A., DeVries, R.P., Deniz, S., Cherian, S., Sunmola, I., Song, X., Marshall, J.J., Gruner, K.A., Daigle, T.L., et al. (2022). Intersectional mapping of multi-transmitter neurons and other cell types in the brain. Cell Rep 40, 111036. 10.1016/j.celrep.2022.111036.

5. Root, D.H., Mejias-Aponte, C.A., Zhang, S., Wang, H.L., Hoffman, A.F., Lupica, C.R., and Morales, M. (2014). Single rodent mesohabenular axons release glutamate and GABA. Nat Neurosci 17, 1543–1551. 10.1038/nn.3823.

6. Meye, F.J., Soiza-Reilly, M., Smit, T., Diana, M.A., Schwarz, M.K., and Mameli, M. (2016). Shifted pallidal co-release of GABA and glutamate in habenula drives cocaine withdrawal and relapse. Nat Neurosci 19, 1019–1024. 10.1038/nn.4334.

7. Lazaridis, I., Tzortzi, O., Weglage, M., Martin, A., Xuan, Y., Parent, M., Johansson, Y., Fuzik, J., Furth, D., Fenno, L.E., et al. (2019). A hypothalamus-habenula circuit controls aversion. Mol Psychiatry 24, 1351–1368. 10.1038/s41380-019-0369-5.

8. Yoo, J.H., Zell, V., Gutierrez-Reed, N., Wu, J., Ressler, R., Shenasa, M.A., Johnson, A.B., Fife, K.H., Faget, L., and Hnasko, T.S. (2016). Ventral tegmental area glutamate neurons co-release GABA and promote positive reinforcement. Nat Commun 7, 13697. 10.1038/ncomms13697.

9. Meye, F.J., Lecca, S., Valentinova, K., and Mameli, M. (2013). Synaptic and cellular profile of neurons in the lateral habenula. Front Hum Neurosci 7, 860. 10.3389/fnhum.2013.00860.

10. Hikosaka, O., Sesack, S.R., Lecourtier, L., and Shepard, P.D. (2008). Habenula: crossroad between the basal ganglia and the limbic system. J Neurosci 28, 11825–11829. 10.1523/JNEUROSCI.3463-08.2008.

11. Proulx, C.D., Hikosaka, O., and Malinow, R. (2014). Reward processing by the lateral habenula in normal and depressive behaviors. Nat Neurosci 17, 1146–1152. 10.1038/nn.3779.

12. Hu, H., Cui, Y., and Yang, Y. (2020). Circuits and functions of the lateral habenula in health and in disease. Nat Rev Neurosci 21, 277–295. 10.1038/s41583-020-0292-4.

13. Matsumoto, M., and Hikosaka, O. (2007). Lateral habenula as a source of negative reward signals in dopamine neurons. Nature 447, 1111–1115. 10.1038/nature05860.

14. Jhou, T.C., Fields, H.L., Baxter, M.G., Saper, C.B., and Holland, P.C. (2009). The rostromedial tegmental nucleus (RMTg), a GABAergic afferent to midbrain dopamine neurons, encodes aversive stimuli and inhibits motor responses. Neuron 61, 786–800. 10.1016/j.neuron.2009.02.001.

15. Jhou, T.C., Geisler, S., Marinelli, M., Degarmo, B.A., and Zahm, D.S. (2009). The mesopontine rostromedial tegmental nucleus: A structure targeted by the lateral habenula that projects to the ventral tegmental area of Tsai and substantia nigra compacta. J Comp Neurol 513, 566–596. 10.1002/cne.21891.

16. Hong, S., Jhou, T.C., Smith, M., Saleem, K.S., and Hikosaka, O. (2011). Negative reward signals from the lateral habenula to dopamine neurons are mediated by rostromedial tegmental nucleus in primates. J Neurosci 31, 11457–11471. 10.1523/JNEUROSCI.1384-11.2011.

17. Bromberg-Martin, E.S., Matsumoto, M., Nakahara, H., and Hikosaka, O. (2010). Multiple timescales of memory in lateral habenula and dopamine neurons. Neuron 67, 499–510. 10.1016/j.neuron.2010.06.031.

18. Ji, H., and Shepard, P.D. (2007). Lateral habenula stimulation inhibits rat midbrain dopamine neurons through a GABA(A) receptor-mediated mechanism. J Neurosci 27, 6923–6930. 10.1523/JNEUROSCI.0958-07.2007.

19. Brown, P.L., Palacorolla, H., Brady, D., Riegger, K., Elmer, G.I., and Shepard, P.D. (2017). Habenula-Induced Inhibition of Midbrain Dopamine Neurons Is Diminished by Lesions of the Rostromedial Tegmental Nucleus. J Neurosci 37, 217–225. 10.1523/JNEUROSCI.1353-16.2016.

20. Stopper, C.M., and Floresco, S.B. (2014). What’s better for me? Fundamental role for lateral habenula in promoting subjective decision biases. Nat Neurosci 17, 33–35. 10.1038/nn.3587.

21. Lee, H., and Hikosaka, O. (2022). Lateral habenula neurons signal step-by-step changes of reward prediction. iScience 25, 105440. 10.1016/j.isci.2022.105440.

22. Tian, J., and Uchida, N. (2015). Habenula Lesions Reveal that Multiple Mechanisms Underlie Dopamine Prediction Errors. Neuron 87, 1304–1316. 10.1016/j.neuron.2015.08.028.

23. Stephenson-Jones, M., Yu, K., Ahrens, S., Tucciarone, J.M., van Huijstee, A.N., Mejia, L.A., Penzo, M.A., Tai, L.H., Wilbrecht, L., and Li, B. (2016). A basal ganglia circuit for evaluating action outcomes. Nature 539, 289–293. 10.1038/nature19845.

24. Hong, S., and Hikosaka, O. (2008). The globus pallidus sends reward-related signals to the lateral habenula. Neuron 60, 720–729. 10.1016/j.neuron.2008.09.035.

25. Bromberg-Martin, E.S., Matsumoto, M., Hong, S., and Hikosaka, O. (2010). A pallidus-habenula-dopamine pathway signals inferred stimulus values. J Neurophysiol 104, 1068–1076. 10.1152/jn.00158.2010.

26. Li, H., Pullmann, D., and Jhou, T.C. (2019). Valence-encoding in the lateral habenula arises from the entopeduncular region. Elife 8. 10.7554/eLife.41223.

27. Locantore, J., Liu, Y., White, J., Wallace, J.B., Beron, C., Kraft, E., Sabatini, B., and Wallace, M. (2025). Mixed representations of choice direction and outcome by GABA/glutamate cotransmitting neurons in the entopeduncular nucleus. Elife 13. 10.7554/eLife.100488.

28. Wallace, M.L., Saunders, A., Huang, K.W., Philson, A.C., Goldman, M., Macosko, E.Z., McCarroll, S.A., and Sabatini, B.L. (2017). Genetically Distinct Parallel Pathways in the Entopeduncular Nucleus for Limbic and Sensorimotor Output of the Basal Ganglia. Neuron 94, 138–152 e135. 10.1016/j.neuron.2017.03.017.

29. Kim, S., Wallace, M.L., El-Rifai, M., Knudsen, A.R., and Sabatini, B.L. (2022). Co-packaging of opposing neurotransmitters in individual synaptic vesicles in the central nervous system. Neuron 110, 1371–1384 e1377. 10.1016/j.neuron.2022.01.007.

30. Stopper, C.M., Tse, M.T.L., Montes, D.R., Wiedman, C.R., and Floresco, S.B. (2014). Overriding phasic dopamine signals redirects action selection during risk/reward decision making. Neuron 84, 177–189. 10.1016/j.neuron.2014.08.033.

31. Dabney, W., Kurth-Nelson, Z., Uchida, N., Starkweather, C.K., Hassabis, D., Munos, R., and Botvinick, M. (2020). A distributional code for value in dopamine-based reinforcement learning. Nature 577, 671–675. 10.1038/s41586-019-1924-6.

32. Romero Pinto, S., and Uchida, N. (2025). Tonic dopamine and biases in value learning linked through a biologically inspired reinforcement learning model. Nat Commun 16, 7529. 10.1038/s41467-025-62280-1.

33. Masset, P., Tano, P., Kim, H.R., Malik, A.N., Pouget, A., and Uchida, N. (2025). Multi-timescale reinforcement learning in the brain. Nature 642, 682–690. 10.1038/s41586-025-08929-9.

34. Lowet, A.S., Zheng, Q., Meng, M., Matias, S., Drugowitsch, J., and Uchida, N. (2025). An opponent striatal circuit for distributional reinforcement learning. Nature 639, 717–726. 10.1038/s41586-024-08488-5.

35. Sousa, M., Bujalski, P., Cruz, B.F., Louie, K., McNamee, D.C., and Paton, J.J. (2025). A multidimensional distributional map of future reward in dopamine neurons. Nature 642, 691–699. 10.1038/s41586-025-09089-6.

36. Day, J.J., Jones, J.L., Wightman, R.M., and Carelli, R.M. (2010). Phasic nucleus accumbens dopamine release encodes effort- and delay-related costs. Biol Psychiatry 68, 306–309. 10.1016/j.biopsych.2010.03.026.

37. Sugam, J.A., Day, J.J., Wightman, R.M., and Carelli, R.M. (2012). Phasic nucleus accumbens dopamine encodes risk-based decision-making behavior. Biol Psychiatry 71, 199–205. 10.1016/j.biopsych.2011.09.029.

38. St Onge, J.R., and Floresco, S.B. (2009). Dopaminergic modulation of risk-based decision making. Neuropsychopharmacology 34, 681–697. 10.1038/npp.2008.121.

39. Webber, H.E., Lopez-Gamundi, P., Stamatovich, S.N., de Wit, H., and Wardle, M.C. (2021). Using pharmacological manipulations to study the role of dopamine in human reward functioning: A review of studies in healthy adults. Neurosci Biobehav Rev 120, 123–158. 10.1016/j.neubiorev.2020.11.004.

40. Floresco, S.B., Tse, M.T., and Ghods-Sharifi, S. (2008). Dopaminergic and glutamatergic regulation of effort- and delay-based decision making. Neuropsychopharmacology 33, 1966–1979. 10.1038/sj.npp.1301565.

41. Shabel, S.J., Proulx, C.D., Trias, A., Murphy, R.T., and Malinow, R. (2012). Input to the lateral habenula from the basal ganglia is excitatory, aversive, and suppressed by serotonin. Neuron 74, 475–481. 10.1016/j.neuron.2012.02.037.

42. Wang, D., Li, Y., Feng, Q., Guo, Q., Zhou, J., and Luo, M. (2017). Learning shapes the aversion and reward responses of lateral habenula neurons. Elife 6. 10.7554/eLife.23045.

43. Matsumoto, M., and Hikosaka, O. (2009). Representation of negative motivational value in the primate lateral habenula. Nat Neurosci 12, 77–84. 10.1038/nn.2233.

44. Bromberg-Martin, E.S., and Hikosaka, O. (2011). Lateral habenula neurons signal errors in the prediction of reward information. Nat Neurosci 14, 1209–1216. 10.1038/nn.2902.

45. Wang, W. (2025). Principles of Machine Learning (Springer).

46. Montague, P.R., Dayan, P., and Sejnowski, T.J. (1996). A framework for mesencephalic dopamine systems based on predictive Hebbian learning. J Neurosci 16, 1936–1947.

47. Sutton, R.S., and Barto, A.G. (1998). Reinforcement Learning: An introduction. (MIT Press).

48. Schultz, W., Dayan, P., and Montague, P.R. (1997). A neural substrate of prediction and reward. Science 275, 1593–1599.

49. Hikosaka, O. (2010). The habenula: from stress evasion to value-based decision-making. Nat Rev Neurosci 11, 503–513. 10.1038/nrn2866.

50. Steinberg, E.E., Keiflin, R., Boivin, J.R., Witten, I.B., Deisseroth, K., and Janak, P.H. (2013). A causal link between prediction errors, dopamine neurons and learning. Nat Neurosci 16, 966–973. 10.1038/nn.3413.

51. Chang, C.Y., Esber, G.R., Marrero-Garcia, Y., Yau, H.J., Bonci, A., and Schoenbaum, G. (2016). Brief optogenetic inhibition of dopamine neurons mimics endogenous negative reward prediction errors. Nat Neurosci 19, 111–116. 10.1038/nn.4191.

52. Schoenbaum, G., Esber, G.R., and Iordanova, M.D. (2013). Dopamine signals mimic reward prediction errors. Nat Neurosci 16, 777–779. 10.1038/nn.3448.

53. Tsai, H.C., Zhang, F., Adamantidis, A., Stuber, G.D., Bonci, A., de Lecea, L., and Deisseroth, K. (2009). Phasic firing in dopaminergic neurons is sufficient for behavioral conditioning. Science 324, 1080–1084. 10.1126/science.1168878.

54. de Jong, J.W., Liang, Y., Verharen, J.P.H., Fraser, K.M., and Lammel, S. (2024). State and rate-of-change encoding in parallel mesoaccumbal dopamine pathways. Nat Neurosci 27, 309–318. 10.1038/s41593-023-01547-6.

55. Harkin, E.F., Lynn, M.B., Payeur, A., Boucher, J.F., Caya-Bissonnette, L., Cyr, D., Stewart, C., Longtin, A., Naud, R., and Beique, J.C. (2023). Temporal derivative computation in the dorsal raphe network revealed by an experimentally driven augmented integrate-and-fire modeling framework. Elife 12. 10.7554/eLife.72951.

56. Lowet, A.S., Zheng, Q., Matias, S., Drugowitsch, J., and Uchida, N. (2020). Distributional Reinforcement Learning in the Brain. Trends Neurosci 43, 980–997. 10.1016/j.tins.2020.09.004.

57. Lecca, S., Pelosi, A., Tchenio, A., Moutkine, I., Lujan, R., Herve, D., and Mameli, M. (2016). Rescue of GABAB and GIRK function in the lateral habenula by protein phosphatase 2A inhibition ameliorates depression-like phenotypes in mice. Nat Med 22, 254–261. 10.1038/nm.4037.

58. Lecca, S., Trusel, M., and Mameli, M. (2017). Footshock-induced plasticity of GABAB signalling in the lateral habenula requires dopamine and glucocorticoid receptors. Synapse 71. 10.1002/syn.21948.

59. Tan, D., Nuno-Perez, A., Mameli, M., and Meye, F.J. (2018). Cocaine withdrawal reduces GABAB R transmission at entopeduncular nucleus - lateral habenula synapses. Eur J Neurosci. 10.1111/ejn.14120.

60. Hashikawa, Y., Hashikawa, K., Rossi, M.A., Basiri, M.L., Liu, Y., Johnston, N.L., Ahmad, O.R., and Stuber, G.D. (2020). Transcriptional and Spatial Resolution of Cell Types in the Mammalian Habenula. Neuron 106, 743–758 e745. 10.1016/j.neuron.2020.03.011.

61. Wallace, M.L., Huang, K.W., Hochbaum, D., Hyun, M., Radeljic, G., and Sabatini, B.L. (2020). Anatomical and single-cell transcriptional profiling of the murine habenular complex. Elife 9. 10.7554/eLife.51271.

62. Parent, M., Levesque, M., and Parent, A. (2001). Two types of projection neurons in the internal pallidum of primates: single-axon tracing and three-dimensional reconstruction. J Comp Neurol 439, 162–175.

63. Yu, Z., Guindani, M., Grieco, S.F., Chen, L., Holmes, T.C., and Xu, X. (2022). Beyond t test and ANOVA: applications of mixed-effects models for more rigorous statistical analysis in neuroscience research. Neuron 110, 21–35. 10.1016/j.neuron.2021.10.030.

64. Tanimoto, Y., Kakinuma, H., Aoki, R., Shiraki, T., Higashijima, S.I., and Okamoto, H. (2024). Transgenic tools targeting the basal ganglia reveal both evolutionary conservation and specialization of neural circuits in zebrafish. Cell Rep 43, 113916. 10.1016/j.celrep.2024.113916.

65. Amo, R., Fredes, F., Kinoshita, M., Aoki, R., Aizawa, H., Agetsuma, M., Aoki, T., Shiraki, T., Kakinuma, H., Matsuda, M., et al. (2014). The habenulo-raphe serotonergic circuit encodes an aversive expectation value essential for adaptive active avoidance of danger. Neuron 84, 1034–1048. 10.1016/j.neuron.2014.10.035.

66. Ramaswamy, M., Cheng, R.K., and Jesuthasan, S. (2020). Identification of GABAergic neurons innervating the zebrafish lateral habenula. Eur J Neurosci 52, 3918–3928. 10.1111/ejn.14843.

67. Turner, K.J., Hawkins, T.A., Yanez, J., Anadon, R., Wilson, S.W., and Folgueira, M. (2016). Afferent Connectivity of the Zebrafish Habenulae. Front Neural Circuits 10, 30. 10.3389/fncir.2016.00030.

68. Stephenson-Jones, M., Ericsson, J., Robertson, B., and Grillner, S. (2012). Evolution of the basal ganglia: dual-output pathways conserved throughout vertebrate phylogeny. J Comp Neurol 520, 2957–2973. 10.1002/cne.23087.

69. Diaz, E., Bravo, D., Rojas, X., and Concha, M.L. (2011). Morphologic and immunohistochemical organization of the human habenular complex. J Comp Neurol 519, 3727–3747. 10.1002/cne.22687.

70. Matsumoto, M., and Hikosaka, O. (2011). Electrical stimulation of the primate lateral habenula suppresses saccadic eye movement through a learning mechanism. PLoS One 6, e26701. 10.1371/journal.pone.0026701.

71. Stamatakis, A.M., and Stuber, G.D. (2012). Activation of lateral habenula inputs to the ventral midbrain promotes behavioral avoidance. Nat Neurosci 15, 1105–1107. 10.1038/nn.3145.

72. Proulx, C.D., Aronson, S., Milivojevic, D., Molina, C., Loi, A., Monk, B., Shabel, S.J., and Malinow, R. (2018). A neural pathway controlling motivation to exert effort. Proc Natl Acad Sci U S A 115, 5792–5797. 10.1073/pnas.1801837115.

73. Congiu, M., Mondoloni, S., Zouridis, I.S., Schmors, L., Lecca, S., Lalive, A.L., Ginggen, K., Deng, F., Berens, P., Paolicelli, R.C., et al. (2023). Plasticity of neuronal dynamics in the lateral habenula for cue-punishment associative learning. Mol Psychiatry 28, 5118–5127. 10.1038/s41380-023-02155-3.

74. Lecca, S., Meye, F.J., Trusel, M., Tchenio, A., Harris, J., Schwarz, M.K., Burdakov, D., Georges, F., and Mameli, M. (2017). Aversive stimuli drive hypothalamus-to-habenula excitation to promote escape behavior. Elife 6. 10.7554/eLife.30697.

75. Quina, L.A., Walker, A., Morton, G., Han, V., and Turner, E.E. (2020). GAD2 Expression Defines a Class of Excitatory Lateral Habenula Neurons in Mice that Project to the Raphe and Pontine Tegmentum. eNeuro 7. 10.1523/ENEURO.0527-19.2020.

76. Brinschwitz, K., Dittgen, A., Madai, V.I., Lommel, R., Geisler, S., and Veh, R.W. (2010). Glutamatergic axons from the lateral habenula mainly terminate on GABAergic neurons of the ventral midbrain. Neuroscience 168, 463–476. 10.1016/j.neuroscience.2010.03.050.

77. Zhang, L., Hernandez, V.S., Swinny, J.D., Verma, A.K., Giesecke, T., Emery, A.C., Mutig, K., Garcia-Segura, L.M., and Eiden, L.E. (2018). A GABAergic cell type in the lateral habenula links hypothalamic homeostatic and midbrain motivation circuits with sex steroid signaling. Transl Psychiatry 8, 50. 10.1038/s41398-018-0099-5.

78. Flanigan, M., Aleyasin, H., Takahashi, A., Golden, S.A., and Russo, S.J. (2017). An emerging role for the lateral habenula in aggressive behavior. Pharmacol Biochem Behav 162, 79–86. 10.1016/j.pbb.2017.05.003.

79. Ceballos, C.C., Ma, L., Qin, M., and Zhong, H. (2024). Widespread co-release of glutamate and GABA throughout the mouse brain. Commun Biol 7, 1502. 10.1038/s42003-024-07198-y.

80. Shabel, S.J., Murphy, R.T., and Malinow, R. (2014). Negative learning bias is associated with risk aversion in a genetic animal model of depression. Front Hum Neurosci 8, 1. 10.3389/fnhum.2014.00001.

81. Lin, W., Xu, J., Zhang, X., and Dolan, R.J. (2025). Habenula-ventral tegmental area functional coupling and risk aversion in humans. Proc Natl Acad Sci U S A 122, e2500815122. 10.1073/pnas.2500815122.

82. Herkenham, M., and Nauta, W.J. (1977). Afferent connections of the habenular nuclei in the rat. A horseradish peroxidase study, with a note on the fiber-of-passage problem. J Comp Neurol 173, 123–146. 10.1002/cne.901730107.

83. Muller, T.H., Butler, J.L., Veselic, S., Miranda, B., Wallis, J.D., Dayan, P., Behrens, T.E.J., Kurth-Nelson, Z., and Kennerley, S.W. (2024). Distributional reinforcement learning in prefrontal cortex. Nat Neurosci 27, 403–408. 10.1038/s41593-023-01535-w.

84. Rink, E., and Wullimann, M.F. (2001). The teleostean (zebrafish) dopaminergic system ascending to the subpallium (striatum) is located in the basal diencephalon (posterior tuberculum). Brain Res 889, 316–330. 10.1016/s0006-8993(00)03174-7.

85. Matsui, H. (2017). Dopamine system, cerebellum, and nucleus ruber in fish and mammals. Dev Growth Differ 59, 219–227. 10.1111/dgd.12357.

86. Grillner, S., Robertson, B., and Stephenson-Jones, M. (2013). The evolutionary origin of the vertebrate basal ganglia and its role in action selection. J Physiol 591, 5425–5431. 10.1113/jphysiol.2012.246660.

87. Berger, B., Gaspar, P., and Verney, C. (1991). Dopaminergic innervation of the cerebral cortex: unexpected differences between rodents and primates. Trends Neurosci 14, 21–27. 10.1016/0166-2236(91)90179-x.

88. Davenport, A.T., Grant, K.A., Szeliga, K.T., Friedman, D.P., and Daunais, J.B. (2014). Standardized method for the harvest of nonhuman primate tissue optimized for multiple modes of analyses. Cell Tissue Bank 15, 99–110. 10.1007/s10561-013-9380-2.

89. Turi, G.F., Li, W.K., Chavlis, S., Pandi, I., O’Hare, J., Priestley, J.B., Grosmark, A.D., Liao, Z., Ladow, M., Zhang, J.F., et al. (2019). Vasoactive Intestinal Polypeptide-Expressing Interneurons in the Hippocampus Support Goal-Oriented Spatial Learning. Neuron 101, 1150–1165 e1158. 10.1016/j.neuron.2019.01.009.

