## Supplementary Figures for "Evolutionary rise of a synaptic mechanism for creating and diversifying key reinforcement signals"

### Supplementary Figure 1

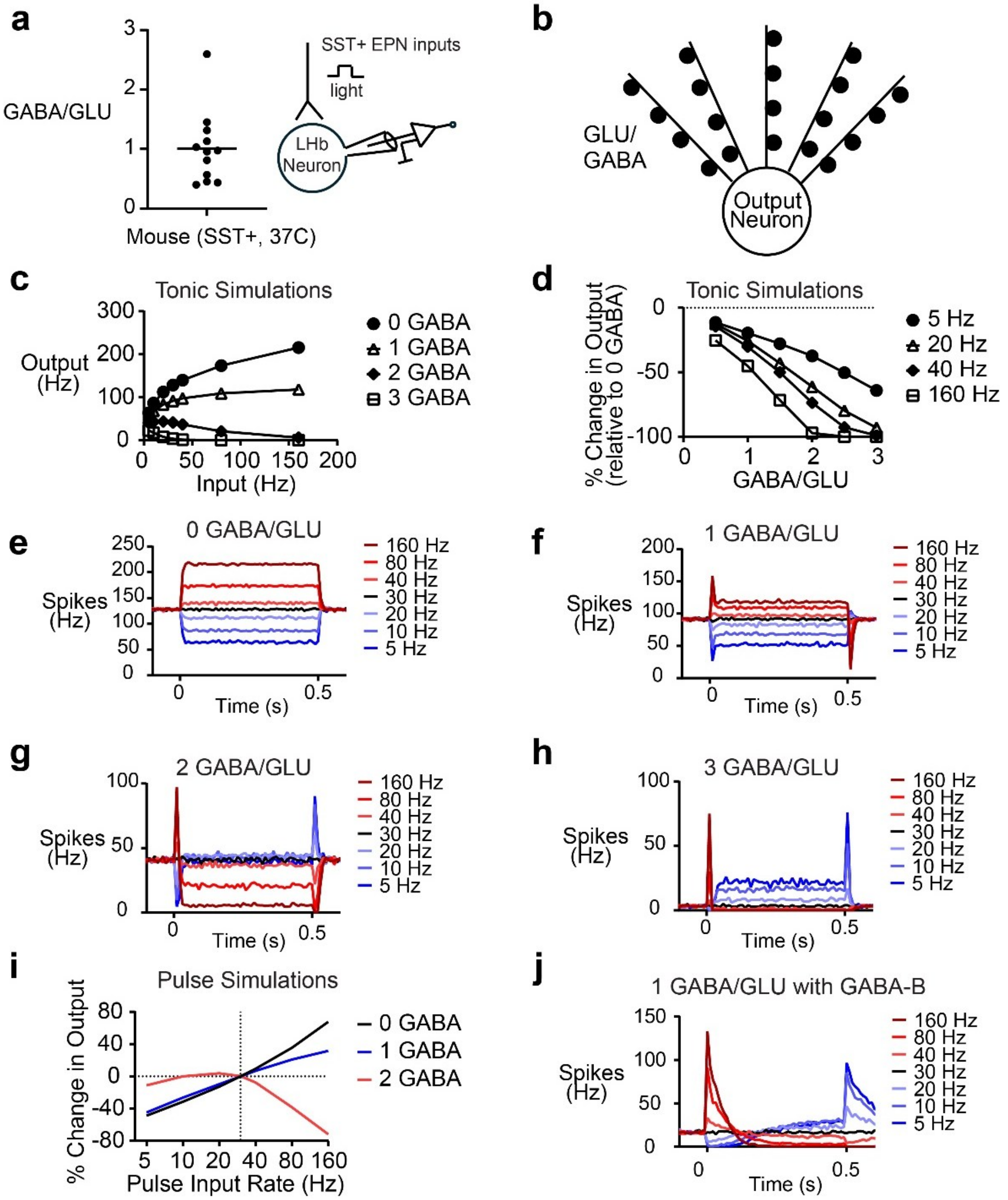


**Fig. S1. Data related to Figure 1. a**, Ratio of peak amplitude of GABA-A-R and AMPA-R-mediated (GLU) currents from whole-cell, ex vivo recordings of mouse LHb neurons (N = 12 neurons from 5 mice; 37 ºC) during optogenetic stimulation of somatostatin-positive (SST+) EPN inputs. **b**, Schematic figure of a neuron used for simulations in **c**-**j** (similar to Figure 1c, except with twice as many input neurons and synapses). **c**, Input-output curves for simulations of tonic input activity with different ratios of GABAergic and glutamatergic peak conductances (average of 10, 3-second simulations; similar to Figure 1e, except with twice as many input neurons and synapses). **d**, Same data as **c**, but recalculated and plotted to show the change in output activity for different amounts of GABA co-transmission (relative to no GABA) and different input activity. **e**-**h**, Output activity during pulse simulations for different amounts of GABA co-transmission, as indicated (using output neuron in **b**). Colors indicate mean firing rate of inputs during the 0.5 s pulse period. **i**, Change in output activity relative to 30 Hz input baseline period for different pulse input rates using output neuron in **b**. **j**, same as **f** but with GABA-B conductance.

#### Supplementary Figure 2

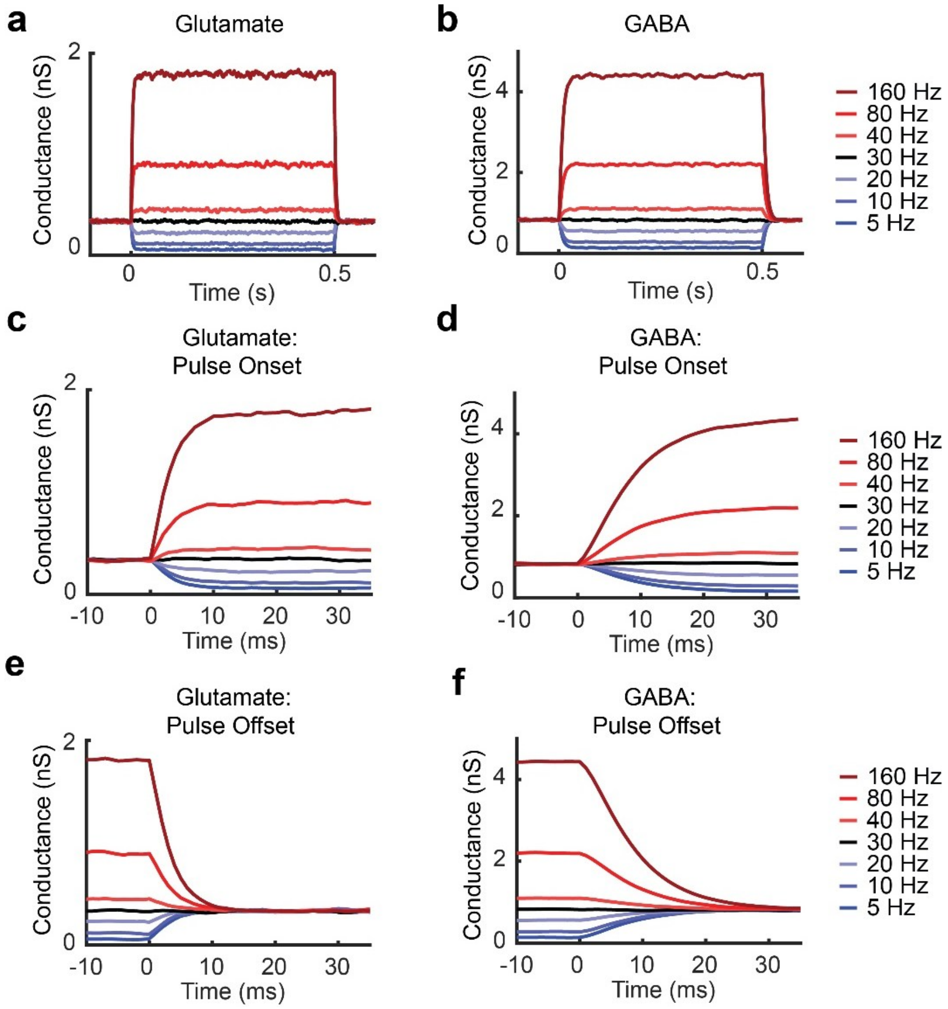


**Fig. S2. Data related to Figure 2.** Mean glutamate (**a**, **c**, **e**) and GABA (**b**, **d**, **f**) conductances at one synapse during pulse simulations (GABA/GLU = 1 condition). Colors indicate input frequency during pulses. Input frequency before pulses is 30 Hz. **c**-**d**, Pulse onset. **e**-**f**, Pulse offset.

### Supplementary Figure 3

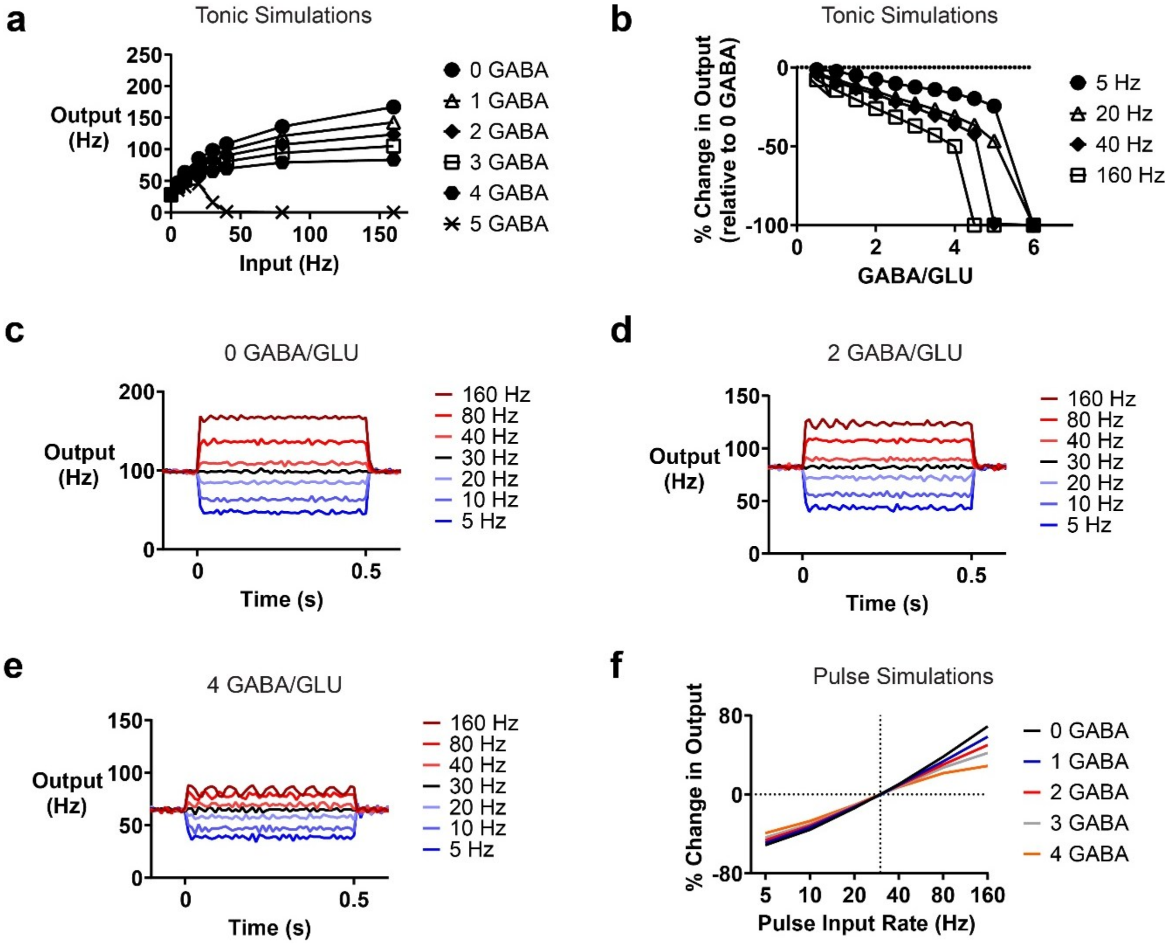


**Fig. S3. Data related to Figure 2. a-f**, same as Figure 2d-i, except that now GABA kinetics are changed to match glutamate kinetics (rather than vice versa). **a**, Input-output curves during tonic input activity simulations with equal glutamate and GABA kinetics (GABA kinetics changed to match glutamate kinetics; average of 10, 3-second simulations). **b**, Same data as **a**, but recalculated and plotted to show the change in output activity for different amounts of GABA co-transmission (relative to no GABA) and different input activity. **c-e**, Output activity for different amounts of GABA co-transmission during pulse simulations with artificially equal glutamate and GABA kinetics (GABA kinetics changed to match glutamate kinetics). **f**, Change in output activity relative to 30 Hz input baseline period for different pulse input rates.

### Supplementary Figure 4

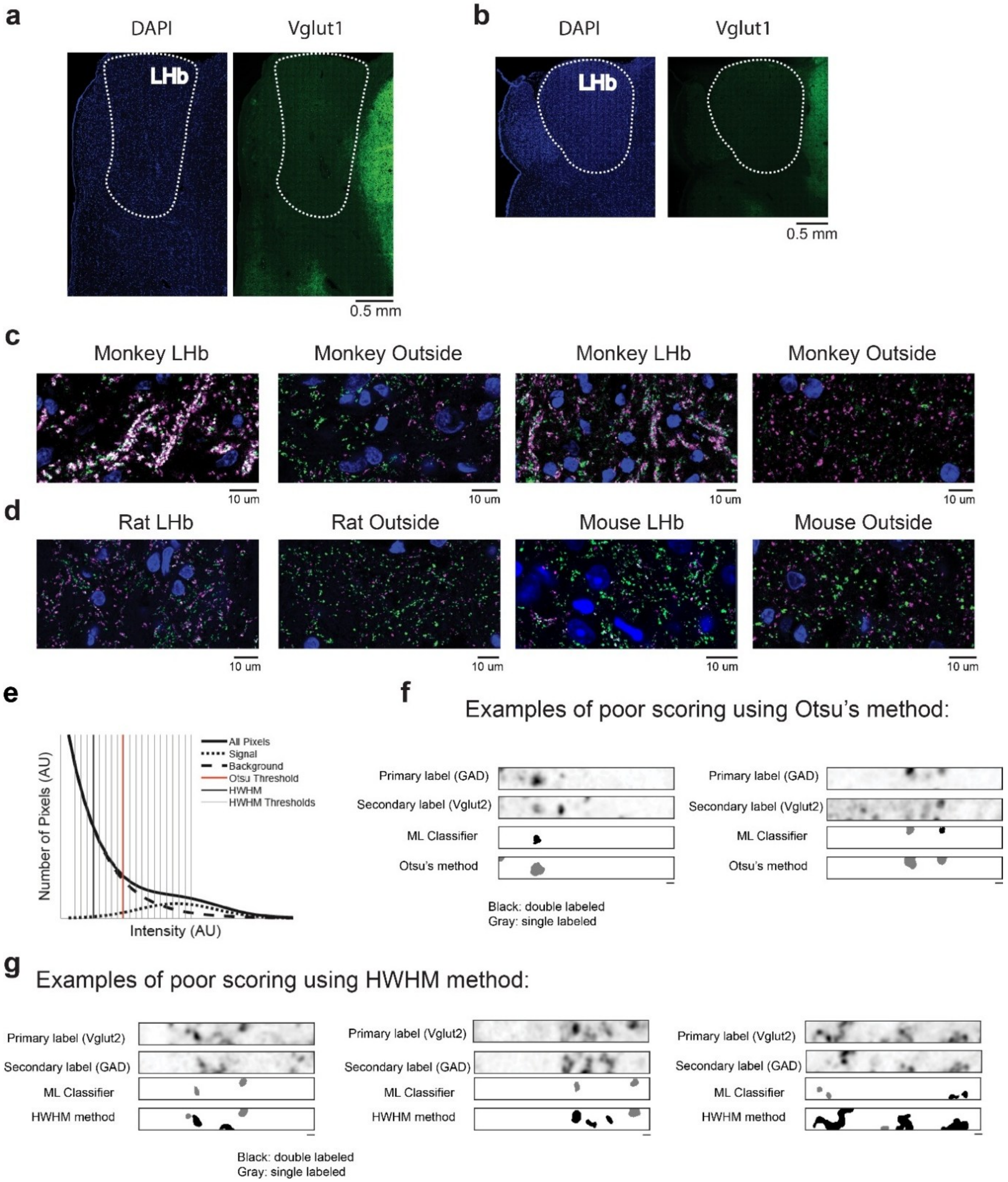


**Fig. S4. Data related to Figure 3. a**-**b**, DAPI and Vglut1 fluorescence in two monkeys. Dashed line indicates LHb. **c**, Examples of "ribbon-like" synaptic terminal structures in the monkey LHb, which were less common outside the LHb. Green, Vglut2. Purple, GAD. Blue, DAPI. **d**, Example synaptic terminals in rodents for comparison with **c**. **e**, Diagram showing Otsu's method and HWHM method for distribution thresholding of background and signal fluorescence. **f**, Examples of poor scoring using Otsu's method for separating background fluorescence from signal. Scale, 1 um. **g**, Examples of poor scoring using the HWHM method for separating background fluorescence from signal. These examples used the best performing HWHM thresholds from the ROC analysis (i.e., largest Youden score). Scale,1 um.

### Supplementary Figure 5

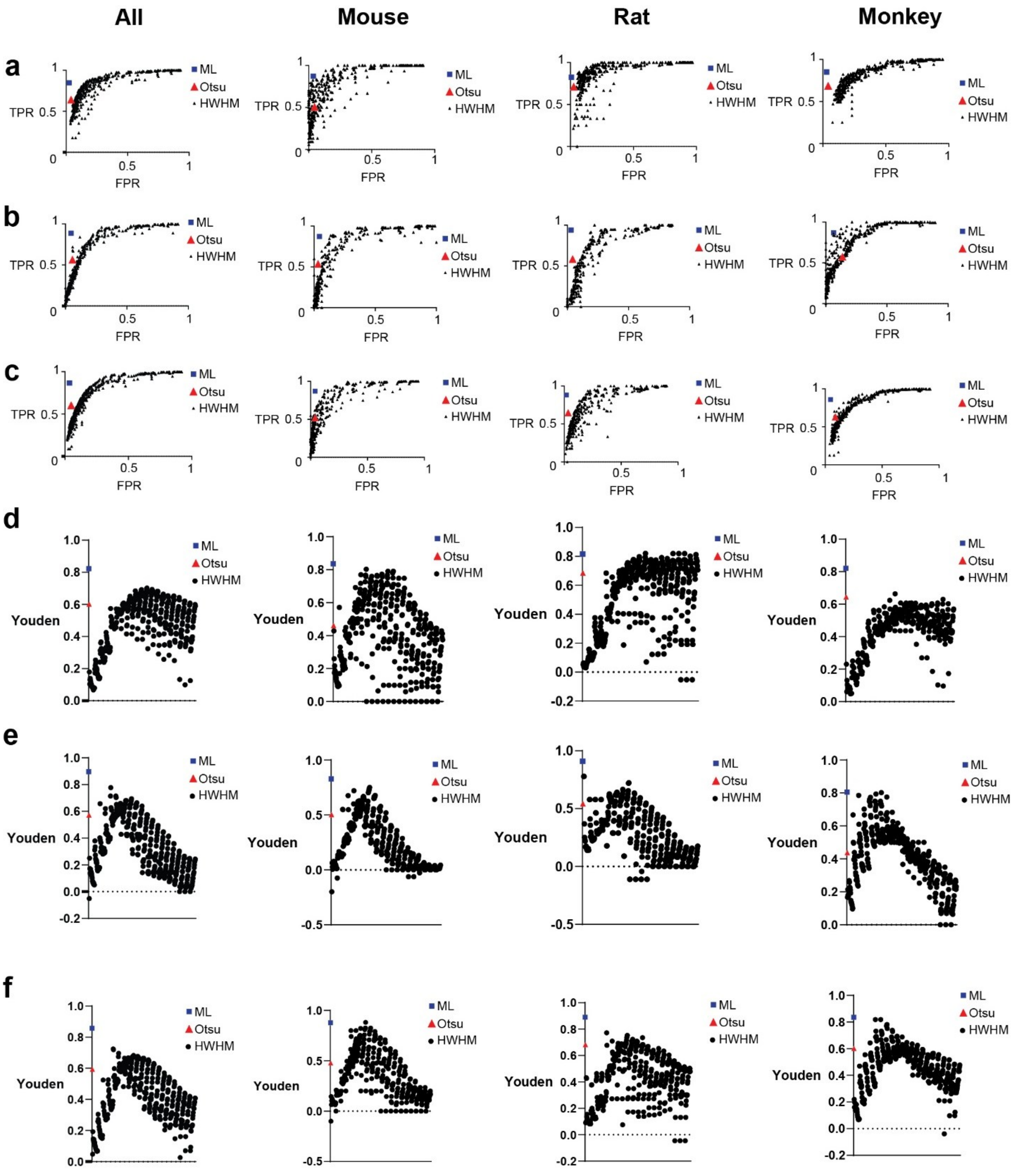


**Fig. S5. Data related to Figure 3.** Column titles (All, Mouse, Rat, Monkey) refer to all data in the figure. **a**, ROC plots for scoring GAD in Vglut2+ terminals in all training images. **b**, Like **a**, but for scoring Vglut2 in GAD+ terminals. **c**, Like **a**, but for average of all scoring. **d**, Youden's index for scoring GAD in Vglut2+ terminals. **e**, Like **d**, but for scoring Vglut2 in GAD+ terminals. **f**, Scoring for images that were not used to train the machine classifier (for plots for all training images, see Figure 3 in the main text).

#### Supplementary Figure 6

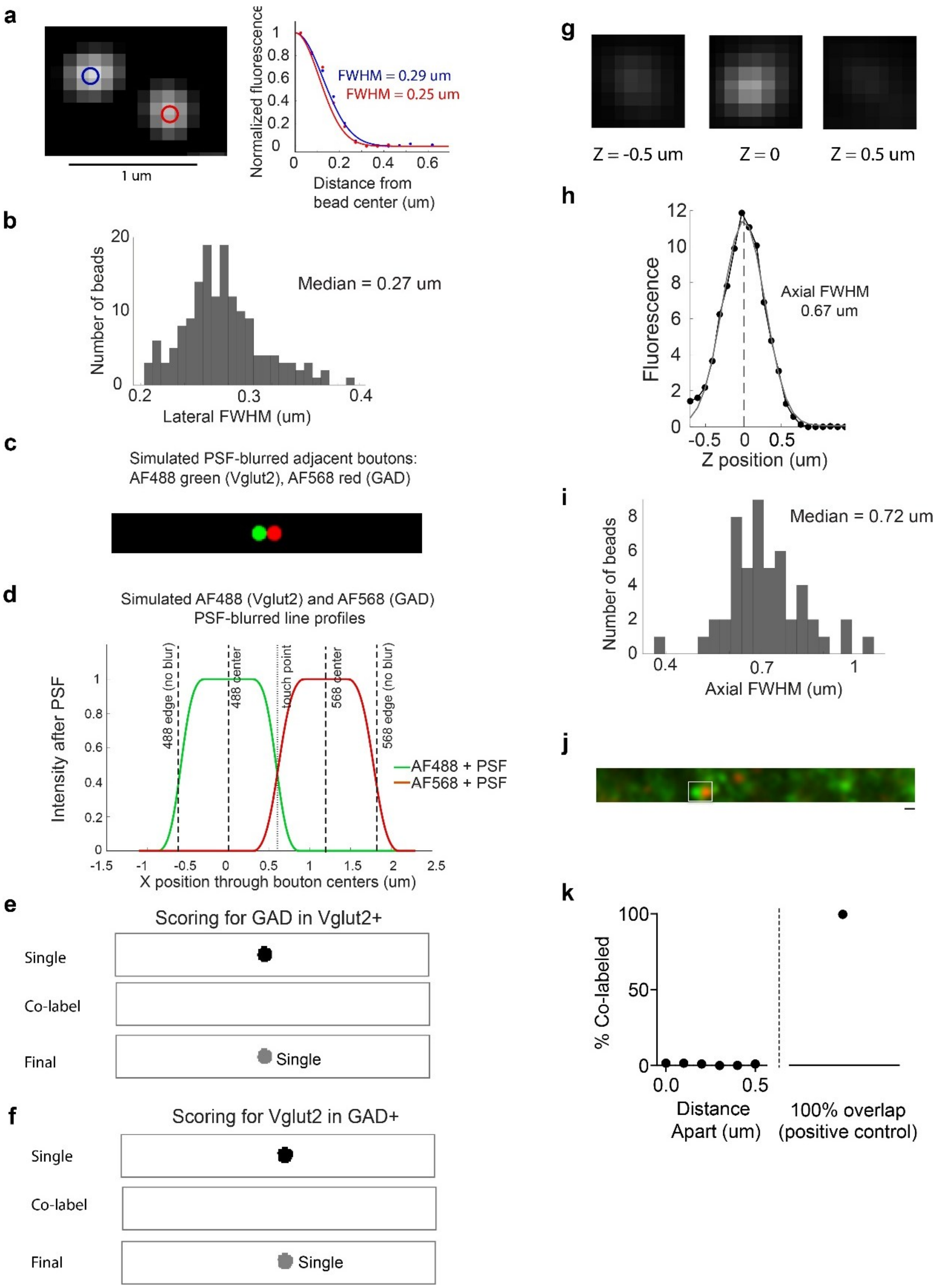


**Fig. S6. Data related to Figure 3. Measurements and tests of the confocal optics. a**, Left: Example fluorescence from two 50 nm green fluorescent beads using the same optics and settings as for our experimental images. Right: Normalized fluorescence relative to the center of each of the beads shown at left. **b**, Histogram of lateral FWHM values from 156 beads. **c**, Simulated fluorescence from immediately laterally adjacent Vglut2+ (Alexa Fluor 488) and GAD+

(Alexa Fluor 568) boutons, blurred according to the empirical lateral FWHM value for the green beads (270 nm) and calculated lateral FWHM for red fluorophores (315 nm). 1.2 um bouton diameter, calculated from the mean LHb bouton volume from Steinkellner et al., *Genetic Probe for Visualizing Glutamatergic Synapses and Vesicles by 3D Electron Microscopy.* ACS Chemical Neuroscience. 2021;12(4):626–639. **d**, Line plot of simulated PSF-blurred fluorescence from c. **e**, GAD (AF568) in Vglut2+ (AF488) scoring of the PSF-blurred simulated boutons in c and d, using our machine learning classifier. No pixels or boutons were misclassified as co-labeled. We found identical results across bouton diameters of 0.4 – 2.0 um. Note that the majority of scored pixels in a bouton would need to be classified as co-labeled for the bouton to be classified as co-labeled. **f**, Same as e, but Vglut2+ (AF488) in GAD (AF568) scoring. **g**, Example fluorescence from one bead shown in different Z (axial) planes. **h**, Fluorescence of same bead in g across different Z planes. **i**, Histogram of Z (axial) FWHM values for 51 beads. **j**, Example image used in the 3D adjacent bouton analysis. PSF-blurred, simulated boutons are outlined by the white rectangle. **k**, Results of the 3D adjacent bouton (mix of 0.8 – 1.6 um bouton diameters) analysis that blurred synthetic adjacent boutons according to our empirical FWHM fluorescence measurements shown in b and i. Only ~ 1% of green boutons were misclassified as co-labeled at 0 um distance (where 0 indicates immediately adjacent spheres).

#### Supplementary Figure 7

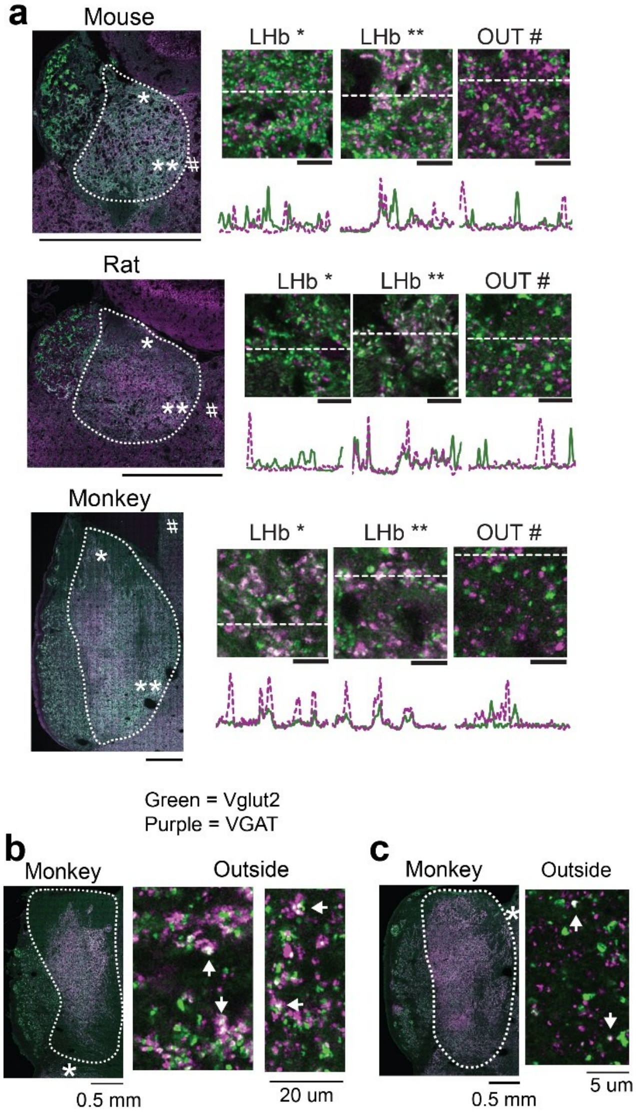


**Fig. S7. Data related to Figure 4. a**, Examples of high-resolution, tiled images from mouse, rat, and monkey, as well as magnified images to the right. Green, Vglut2. Purple, VGAT. White, co-labeling. Scale bars and line plots as in Figure 3a-f. **b**-**c**, Examples of Vglut2 and GAD co-labeling (white) outside of the habenula in monkeys. Dashed line, LHb. * corresponds to location of more magnified images.

#### Supplementary Figure 8

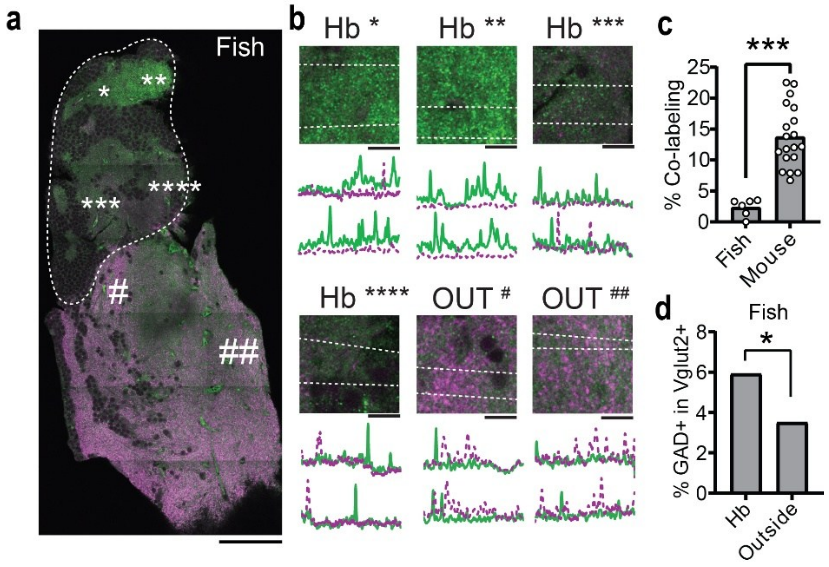


**Fig. S8. Data related to Figure 4. a**, Example high-resolution, tiled image from zebrafish. Green, Vglut2. Purple, GAD, Dashed line, habenula. Scale, 100 µm. *,**,***,****, #, and ## correspond to images in **b**. **b**, Magnified images from corresponding placements in **a**. Line plots as in Figure 3. Scale, 10 µm. **c**, Comparison of co-labeling of Vglut2 and GAD in synaptic terminals from the zebrafish habenula and mouse LHb. **d**, Percentage of glutamatergic terminals that contain GAD in the zebrafish habenula (Hb; N = 354 Vglut2+ terminals) and surrounding regions (Outside; 1051 Vglut2+ terminals), P = .049, chi-square. For **c,d**: *, P < .05; ***, P < .001.

### Supplementary Figure 9

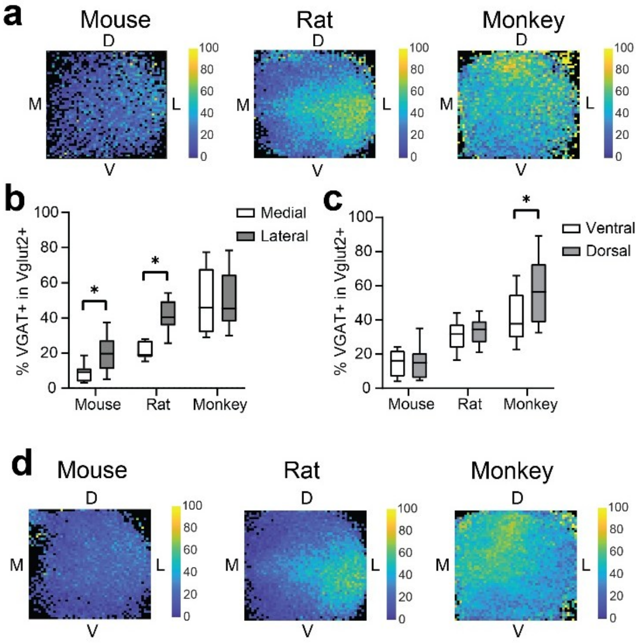


**Fig. S9. Data related to Figure 6. a**, Topographical maps of the percentage of glutamatergic terminals that express VGAT in the mouse (average of 14 tiled images), rat (average of 14 tiled images), and monkey (average of 13 tiled images) LHb. Black indicates no data for the particular tile. **b**, Percentage of glutamatergic terminals that express VGAT in the medial LHb and lateral LHb for each species (same tiled images as in **a**). **c**, same as **b**, but for the dorsal and ventral LHb. **d**, Topographical map of the percentage of glutamatergic terminals that contain GAD in the mouse, rat, and monkey LHb. These are the same as Figure 6d-f, except without the minimum number of slices requirement (i.e., black indicates no data whatsoever). * in **b** and **c**, $P < .0001$.

#### Supplementary Figure 10

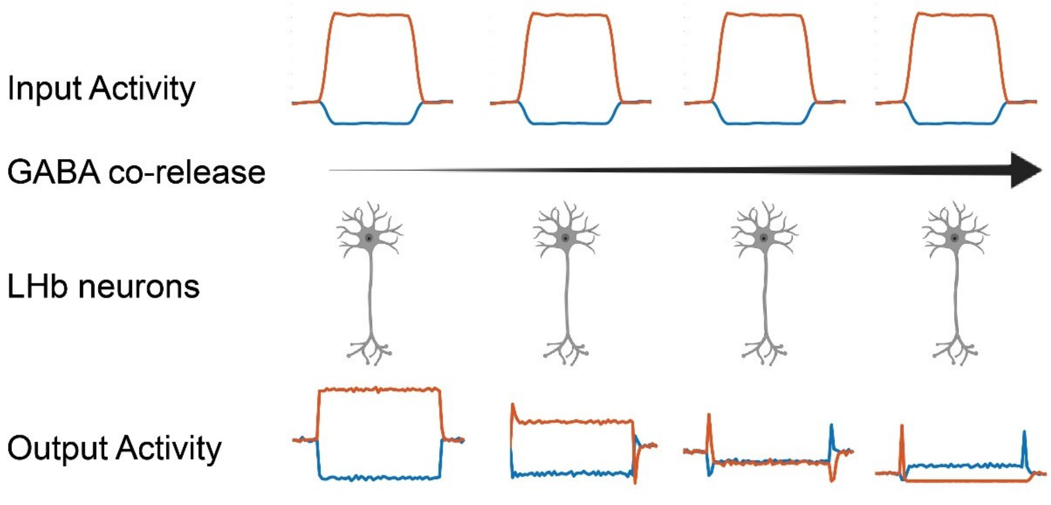


**Fig. S10.** Model of how GABA co-release with glutamate affects input activity transformations in the LHb. The input activity shown is not designed to replicate experimental results but to show how LHb neurons would respond to sudden increases and decreases in input activity.
